# Land-use planning for health: Tradeoffs and nonlinearities govern how land-use change impacts vector-borne disease risk

**DOI:** 10.1101/2021.06.09.447801

**Authors:** Morgan P Kain, Andrew J MacDonald, Erin A Mordecai, Lisa Mandle

**Affiliations:** Department of Biology, Stanford University, Stanford, CA, 94305, USA; Natural Capital Project, Woods Institute for the Environment, Stanford University, Stanford, CA 94305, USA; Earth Research Institute, University of California Santa Barbara, Santa Barbara, CA 93106, USA

**Keywords:** **Keywords**: Ecosystem Services, Land-sparing, Land-sharing, *Nyssorhynchus darling*, R_0_

## Abstract

Patterns of land-use can affect the transmission of many infectious diseases with human health implications; yet, applied ecosystem service models have rarely accounted for disease transmission risk. A mechanistic understanding of how land-use changes alter infectious disease transmission would help to target public health interventions and to minimize human risk of disease with either ecosystem degradation or restoration. Here, we present a spatially explicit model of disease transmission on heterogeneous landscapes that is designed to serve as a road map for modeling the multifaceted impacts of land-use on disease transmission. We model the transmission of three vector-borne diseases with distinct transmission dynamics (parameterized using published literature to represent dengue, yellow fever, and malaria) on simulated landscapes of varying spatial heterogeneity in tree cover and urban area. Overall, we find that these three diseases depend on the biophysical landscape in different nonlinear ways, leading to tradeoffs in disease risk across the landscape; rarely do we predict disease risk to be high for all three diseases in a local setting. We predict that dengue risk peaks in areas of high urban intensity and human population density, yellow fever risk peaks in areas with low to moderate human population density and high tree cover, and malaria risk peaks where patchy tree cover abuts urban area. To examine how this approach can inform land use planning, we applied the model to a small landscape to the northwest of Bogotá, Colombia under multiple restoration scenarios. We predict that in an area inhabited by both *Aedes aegypti* and *Ae. albopictus*, any increase in overall tree cover would increase dengue and yellow fever risk, but that risk can be minimized by pursuing a large contiguous reforestation project as opposed to many small, patchy projects. A large contiguous reforestation project is also able to both reduce overall malaria risk and the number of malaria hotspots. As sustainable development goals make ecosystem restoration and biodiversity conservation top priorities, it is imperative that land use planning account for potential impacts on both disease transmission and other ecosystem services.

**Open Research statement:** All data and code used in this study are available in the online supplemental material. Code and data are also hosted at: https://github.com/morgankain/Land-Use_Disease_Model.

## Introduction

Land-use change, such as the conversion of forest to cropland, rangeland, or urban area, often has negative and long-lasting (or irreversible) impacts on biodiversity (Huston, 2005, Mattison and Norris, 2005, Hansen et al., 2012, Cunningham et al., 2013) and ecosystem services (e.g., water quality: Ren et al. 2003; carbon sequestration: Guo and Gifford 2002; nitrogen cycling and soil quality: Mirza et al. 2014). Though tradeoffs between economic productivity (e.g., crop yield) and environmental values are sometimes inevitable, careful *a priori* planning can help to reduce the severity of these trade-offs and thus minimize environmental degradation (Green et al., 2005, Polasky et al., 2008, Nelson et al., 2009, Carreñ o et al., 2012, Goldstein et al., 2012, Kennedy et al., 2016, Pennington et al., 2017). For example, alternative land-use change scenarios can be compared using an optimization framework where the economic driver of the land-use change (e.g., crop yield), biodiversity, and ecosystem services (such as water quality, carbon sequestration, nutrient retention, and recreation opportunities) are estimated for each scenario and plotted against each other relative to a hypothetically achievable “efficiency frontier” (e.g., see Pennington et al., 2017). The search for optimal land-allocation decisions (e.g., location, patch size, and configuration) using this method is increasing in applied research and contributing to policy decisions (Viglizzo and Frank, 2006, Goldstein et al., 2012, Geneletti, 2013, Kennedy et al., 2016, Pennington et al., 2017).

Land-use change can also affect the transmission of infectious diseases, which are one of the leading global health burdens, as measured by lost disability adjusted life years (DALYs). In 2018, malaria alone accounted for over 228 million cases (with approximately 1/3 - 1 DALY lost per case: Abdalla et al. 2007, Gunda et al. 2016) and an estimated 405,000 deaths (WHO, 2019). Land-use change such as deforestation can increase malaria transmission (Coluzzi, 1994, Sharma, 2002, Hahn et al., 2014), while landscape fragmentation increases human risk of Lyme disease (Ward and Brown, 2004), and dengue transmission tends to increase with urbanization (Vanwambeke et al., 2007). Despite this longstanding knowledge, the health burden of infectious diseases, and decades-old suggestions that infectious disease transmission should be considered when making land-use decisions (e.g., Patz et al., 2004), disease transmission has rarely been accounted for explicitly in applied ecosystem services research. Recently, Castro et al. (2019) provided guidelines for how planning decisions in the Amazon basin could consider infectious disease transmission (with a focus on malaria and dengue) and McClure et al. (2019) identified land-use change-mediated alterations to human population density, human-wildlife contact rates, vector abundance, and human exposure to vectors as the primary mechanisms altering infectious disease transmission. While these works will hopefully bring increased attention to infectious disease transmission in ecosystem services research, neither provided direct advice on how to quantify disease transmission as a function of land-use change. Given the high rates of land-use change globally (e.g., deforestation: Runyan and D’Odorico 2016; urbanization: Seto et al. 2013) and the growing momentum of efforts to accelerate the pace of restoration and protection globally (e.g., the “UN decade on Ecosystem Restoration”: UN 2019; the Global Deal For Nature: Dinerstein et al. 2019), it is a critical time to seek a more quantitative understanding of how land-use changes will impact infectious disease transmission.

How should infectious disease transmission be modeled in order to optimize land-use decisions? Ideally, a model would have three characteristics. First, it should be constructed using a mechanistic understanding of disease dynamics in order to estimate *a priori* how alternative proposed land-use scenarios would affect transmission. Previous research has investigated links between land-use characteristics and disease transmission in a variety of disease systems (e.g., malaria: Vittor et al. 2006, Chaves et al. 2018, San- tos and Almeida 2018, MacDonald and Mordecai 2019; dengue: Ziemann et al. 2018; Lyme disease: Jackson et al. 2006, MacDonald et al. 2019; multiple diseases: Vanwambeke et al. 2007, Sheela et al. 2017); however, these works rely primarily on regression frameworks that are not appropriate for predicting responses to new environmental regimes that have not yet been observed, which is needed to evaluate alternative potential land-use scenarios. Second, a suitable model should be able to link spatially-explicit land-use patterns to disease transmission in order to understand which populations would experience the highest infection risk for planning targeted control efforts and to map tradeoffs or synergies between disease transmission and other important outcomes (e.g., ecosystem services). Finally, flexible, generalizable model frameworks can allow us to compare land use change impacts on multiple diseases, and to capture tradeoffs among them.

Because different infectious diseases often have strikingly different transmission dynamics, predicting the effects of land-use on the transmission of any single disease could require its own modeling study. For example, both cholera, caused by a bacteria passed from an infected to susceptible person through fecal contamination of water, and malaria, a *Plasmodium* parasite transmitted between humans by *Anopheles* (*Nyssorhynchus*) mosquito vectors, may respond to landscape features such as the location, density, and size of human settlements, water bodies, and forest patches. However, because cholera is environmentally transmitted and malaria vector transmitted, the data and modeling techniques required to predict how land-use change would affect the transmission of these two diseases will differ. Even within a smaller class of diseases such as mosquito-borne diseases, the disease-causing pathogens vary in the number of species of hosts and mosquito they use for transmission (altering how humans become infected), which changes the data requirements to model transmission and can promote alternative modeling strategies. For example, Zika virus exploits humans as its primary host and is transmitted primarily by *Aedes aegypti* mosquitoes, which allows single-host single-vector differential equation models to have moderate success (e.g., Bonyah et al., 2017, Riou et al., 2017). On the other hand, Ross River virus circulates in dozens of vertebrate host and mosquito species (Stephenson et al., 2018), which requires data on a much broader array of species in order to parameterize a mechanistic model (e.g., Kain et al., 2021). Further, because the host and vector community of any vector-borne disease will vary across space and time, modeling even a single disease requires, at a minimum, location-specific parameter values; transmission predictions in one community are unlikely to translate well to a different community. The dominant agent of malaria in most of Latin America (*Plasmodium vivax*), for example, is transmitted by *Nyssorhynchus darlingi* (previously *Anopheles darlingi*), which breeds in standing water at forest boundaries (Tadei et al., 1998, Vittor et al., 2006, Zeilhofer et al., 2007, Vittor et al., 2009), whereas the dominant agent of malaria in sub-Saharan Africa (*Plasmodium falciparum*) is transmitted primarily by *Anopheles gambiae*, which readily breeds in urban and peri-urban areas in artificial containers, roadside ditches, and a variety of other small water bodies (Minakawa et al., 2004, Awolola et al., 2007, Gnémé et al., 2019).

A single spatially explicit mechanistic model will not be able to make predictions for a wide variety of diseases given that each disease depends on disease-by-location-specific parameter values. However, general modeling scaffolds exist to quantify the transmission of a broad range of infectious diseases using a common strategy. Once established, such a model framework can be modified and parameterized as needed to capture the transmission dynamics of the most relevant infectious diseases in a given location. Here we construct transmission matrices composed of the transmission rates between all pairs of species that participate in transmission (which can be parameterized using both laboratory infection data and field data on contact rates among species). We make these transmission matrices spatially dependent by modeling species contact rates as a function of landscape features. Explicitly accounting for the transmission rates between all pairs of species can be data intensive, but has the advantage of being able to flexibly model diseases with a variety of transmission modes including directly transmitted diseases (e.g., chickenpox: Ogunjimi et al., 2009), environmentally transmitted diseases (e.g., chronic wasting disease: Jennelle et al., 2014, Samuel and Storm, 2016), and vector-borne diseases with all forms of transmission strategies (including host-to-vector and vector-to-host transmission: Dobson 2004, as well as vector-to-vector vertical transmission: Lequime and Lambrechts 2014; and direct host-to-host transmission, which can occur for some vector-borne pathogens such as Rift Valley fever virus and Zika virus: Wichgers Schreur et al. 2016, D’Ortenzio et al. 2016).

We use the spatially explicit model to capture the transmission of several vector-borne diseases. We analyze patterns of vector-borne disease risk as a function of land-use in two stages. First, we explore general patterns of vector-borne disease transmission as a function of landscape features using simulated landscapes in order to better isolate how landscape features drive disease risk. For this stage we simulate landscapes of tree cover and urban area (i.e., percent impermeable surface, which we will refer to as urban “intensity”). These landscapes are simulated to vary along a gradient of spatial heterogeneity from one extreme of highly segregated landscape features to one of highly integrated features; along this continuum the average of each landscape feature is constrained to remain the same. This continuum is akin to the “land-sparing” vs. “land-sharing” dichotomy, which draws a comparison between a land allocation strategy that separates or integrates production and conservation, respectively (Green et al., 2005, Phalan et al., 2011, Tscharntke et al., 2012). This paradigm has been used extensively to compare methods for conserving biodiversity (Phalan et al., 2011, Melo et al., 2013, Edwards et al., 2014, Goulart et al., 2016), but has not yet been applied to human health and vector-borne disease transmission, despite its utility (see McClure et al., 2019, Table 1 for a list of studies that have used this conceptual paradigm). For each simulated level of spatial heterogeneity we also predict how disease risk will change depending on human population density in an effort to separate the effects of the spatial configuration of urban intensity from human density. Second, we apply the model in a specific case study to predict how potential vector-borne disease risk would change with alternative strategies of partial reforestation on a region of mixed urban area and farmland northwest of Bogotá Colombia, once again using a range from extremes of “land-sparing” to “land-sharing”. This region has been identified as being within a broader area of high restoration importance (Strassburg et al., 2020), and has recent curated MODIS vegetation indices (e.g., NDVI: Normalized Difference Vegetation Index) available (Gerard et al., 2020). Further, this is a high-elevation region that may experience a higher overall vector-borne disease burden with climate warming (Ryan et al., 2019, Mordecai et al., 2020)

In both stages we model the transmission of three mosquito-borne diseases that differ in which landscape features promote transmission, in order to capture the types of tradeoffs that are likely to occur on real landscapes. Specifically, we model the transmission of: (i) a human-to-human specialist that is transmitted by urban-dwelling mosquitoes (e.g., dengue), (ii) a human-to-human specialist that is instead transmitted by mosquitoes that prefer non-built environments (e.g., malaria), and (iii) a disease that both spills over from non-human hosts but also has the potential for human-to-mosquito-to-human transmission (e.g., yellow fever). We use published literature to parameterize each transmission model for dengue transmitted by *Ae. aegypti* and *Ae. albopictus*, malaria transmitted by *Ny. darlingi*, and yellow fever transmitted by *Ae. aegypti*, *Ae. albopictus*, and *Haemagogus* spp. We present our results as applying to the transmission of dengue, malaria, and yellow fever for brevity, though we suggest caution when interpreting our results as direct estimates for these three diseases because of a relatively poorly defined quantitative relationship between many components of each disease’s transmission and land-use. For the first stage of analysis we examine how the configuration of landscape features (urban intensity and tree cover) and absolute abundance of humans alters human infection risk for diseases with these transmission modes and explore tradeoffs in risk among them. For the second stage of analysis we focus on what strategy of landscape restoration minimizes post-restoration disease risk. In an effort to guide future research, we also identify key empirical gaps in the understanding of the underlying mechanistic disease transmission pathway that strongly affect predictions, as well as suggest strategies for better integration of disease transmission into ecosystem services research in the future.

## Methods

### Model Overview

#### Model outcomes

We calculated two summary metrics of disease transmission on each landscape in order to map disease metrics spatially, both of which rely on R_0_, which describes the number of new infections a single source infection would generate in an otherwise susceptible population. The first metric is R_0_ itself, which we calculated for each landscape cell (computationally, each cell [i, j] of a matrix) assuming that an infected individual were to appear in that cell at the beginning of their infectious period. Thus, R_0_ measures the epidemic potential of a disease if it were to emerge in a specific location on a landscape and can be thought of as the potential of that landscape region to serve as a source of infection for the wider region. R_0_ has a storied history of providing sufficient criteria for affecting change (e.g., the R_0_ of malaria was used to identify sufficient vector control for the completion of the Panama Canal: Coleman-Jones 1999), but also has some known drawbacks as an epidemic metric, including assuming a fully susceptible population with no heterogeneity in transmission (within classes, i.e., host species) and a temporally constant environment. The second metric, which we call force of infection (FOI), also depends on a calculation of R_0_, but summarizes transmission in the opposite direction by quantifying the flow of infection into a given location. Akin but not identical to, the classic definition of FOI (the rate at which susceptible individuals acquire an infectious disease), here we use FOI to describe the conversion of susceptible to infected individuals cell by cell after one infection generation. Thus, FOI can be interpreted as a measure of the overall infection burden a specific landscape region would experience from transmission on the broader landscape. We calculated FOI by summing the number of new human cases a given location on the landscape would experience if infection were to arise in all possible locations on the landscape.

To calculate R_0_ in a given landscape cell for the two human-to-human specialists (parameterized to represent dengue and malaria), we assumed a single human infection appeared in that cell. For these diseases we calculated R_0_ as the total number of second generation human infections generated from the source human infection. To calculate FOI in cell [i, j] for these diseases (which we write as FOI_h_ for FOI on humans) we summed the number of second generation human infections in cell [i, j] across all possible disease emergence locations (all landscape cells). For the disease capable of being transmitted by humans and non-human hosts (yellow fever) we calculated both R_0_ and FOI arising from either a source human infection or a source non-human primate infection. To explore the multiple transmission pathways of this disease, our model separates each R_0_ and FOI value into its component parts, i.e., the number of second generation human and non-human primate infections arising from a source non-human primate infection (the sum of which would be the overall R_0_ attributable to a starting non-human primate infection). We note that these are R_0_*-like* quantities as they are not calculated using the traditional dominant eigenvalue of the pairwise transmission matrix (Diekmann et al., 2010) (which gives the expected number of secondary cases in a heterogeneous community arising from a *typical* infection).

#### Model structure

To better characterize the spatial dependence of R_0_ and FOI of these diseases (for efficiency we will henceforth refer to these three diseases simply as dengue, malaria, and yellow fever) on the landscape and to evaluate the role of different mosquito vectors, we separated transmission into two components: host-to-mosquito transmission, which considers the number of mosquitoes the source host infects, and mosquito-to-host transmission, which considers the number of new (second generation) host infections generated from the mosquitoes infected in the host-to-mosquito transmission step. We first describe the overall calculation of host-to-mosquito and mosquito-to-host transmission and then describe in detail the data and statistical models used to parameterize each of the components of these calculations. Figure 1 provides a visual model schematic meant to aid the interpretation of the equations for host-to-mosquito (Eq. 1) and mosquito-to-host (Eq. 2) transmission.

**Figure 1:**
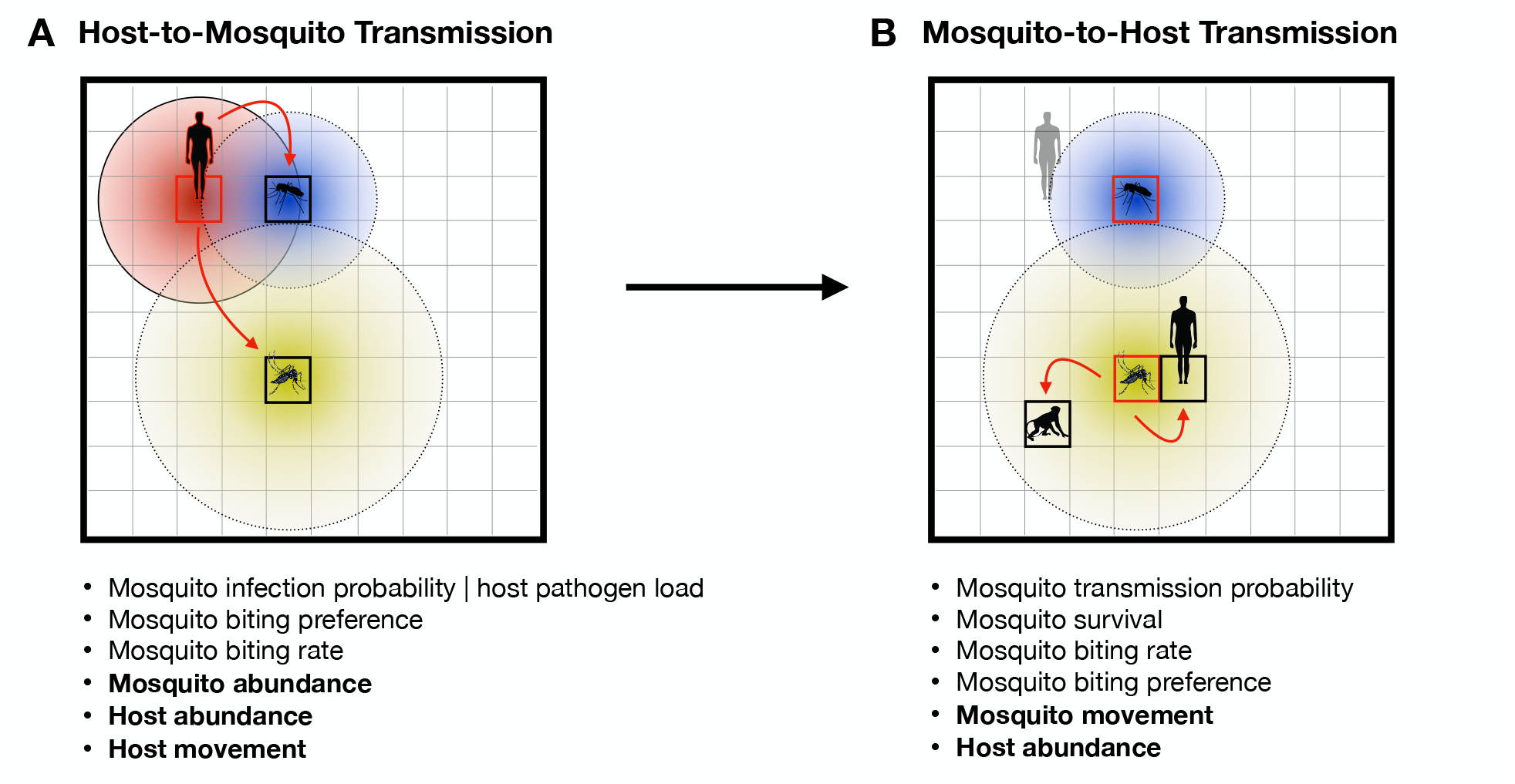
The transmission of a mosquito-borne disease on a landscape starting from a source infection in a human. Red outlined hosts and mosquitoes designate infected individuals, solid black/black outlined individuals are susceptible, and the faded host is recovered. Red arrows show transfer of infection from infected to susceptible individuals. Colored circles in **A** and **B** represent the movement distributions of hosts and mosquitoes on the gridded landscape; boxed cells show the center of these distributions. The text lists the components considered in each transmission step; components that are bolded depend on landscape features. We use the total number of second generation human infections (solid human receiving infection from the red boxed mosquito in **B**) as the R_0_ of an infection originating in the boxed cell in **A**. For dengue and malaria, blood feeding by the infected mosquito on non-human hosts (e.g., the non-human primate in **B**) do not contribute to onward transmission (but do so for yellow fever). Force of infection looks at the problem in the opposite direction, measuring the burden each cell experiences from infections beginning in all possible cells (for which the boxed cell in **A** is one possibility). That is, the transmission pathway pictured here provides one entry in the calculation of FOI for each landscape cell.

All of the mosquitoes across the landscape that become infected by a single infection emerging in landscape cell [i, j] (e.g., the red boxed cell in Figure 1A) can be written as:

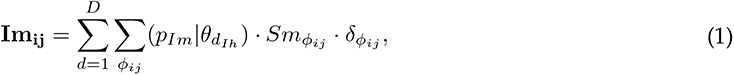

where *h* and *m* refer to hosts and mosquitoes, *I* and *S* refer to infected and susceptible individuals, and terms with the subscripts *ij* designate specific landscape cells. For ease of interpretation we write this equation for one susceptible mosquito species; multiple species could be represented with an additional subscript on each term. The outcome matrix **Im_ij_** contains the total number of mosquitoes with a home landscape cell [i, j] (the center of a mosquito’s flight distribution, see Figure 1A) that get infected by feeding on the source infected host as it moves about the landscape (in Figure 1A, cells within the infected host’s red circle). We use *φ_ij_* to represent the cells that the infected host enters. The total number of mosquitoes that become infected (Figure 1A) is given by a pair of sums: a sum over the host’s movement on the landscape (sum over *φ_ij_*), and a sum over the host’s infectious period (sum over *d*). Internal to these two sums are three terms that define host-to-mosquito transmission within the *φ_ij_* cells: (*p_Im_|θ_dIh_*), the probability of host-to-mosquito transmission during blood feeding; *Sm_φij_*, the number of susceptible mosquitoes in cell *φ_ij_* ; and *δ_φij_*, the biting rate of each mosquito on the infected host in cell *φ_ij_* . We now unpack these three terms. Susceptible mosquitoes can become infected by feeding on the infected host within any cell *φ_ij_* that is found within the mosquitoes’ flight distribution (e.g., in Figure 1A both the blue mosquito’s distribution and the yellow mosquito’s distribution overlap some portion of the infected host’s red distribution). A greater overlap in the host’s and mosquito’s distributions increases their contact rate, leading to more opportunities for blood feeding and thus transmission. While the infected host is in a given cell *φ_ij_*, a susceptible mosquito can, but is not guaranteed to, become infected during a blood feeding event on the infected host. The transmission probability during a feeding event is a function of the host’s pathogen load (which varies over the course of their infection) and the intrinsic ability of the mosquito species to become infected. We write this as (*p_Im_|θ_dIh_*), where *θ_dIh_* describes the host’s pathogen load on day *d* of their infectious period.

The total number of feeding events (and thus potential mosquito infections) that occur while the host is in cell *φ_ij_* is first an increasing function of the total number of mosquitoes in that cell. We use *Sm_φij_* to designate the number of susceptible mosquitoes in cell *φ_ij_*, which is a function of the landscape suitability in and around *φ_ij_* . We note that in the model we keep track of the home cell origin of each of the mosquitoes that make up *Sm_φij_* in order to accurately calculate **Im_ij_**; however, we write *Sm_φij_* as a total number of mosquitoes here for simplicity instead of including additional subscripts designating the home origin of each of these mosquitoes. The total number of feeding events is also a function of the feeding rate of each mosquito on the infected individual, which is a decreasing function of the number of susceptible hosts which pull bites away from the infected host; an increasing function of the mosquitoes’ intrinsic feeding preference on the species identity of the infected host; and finally an increasing function of mosquitoes’ general feeding rate. We collapse these three terms into *δ_φij_*, but detail each of these terms, as well as all of the other Eq. 1 components after Eq. 2.

All of the second generation hosts across the landscape that become infected (**Ih_ij_**) by the mosquitoes calculated in the host-to-mosquito transmission step (**Im_ij_**; Eq. 1), can be written as:

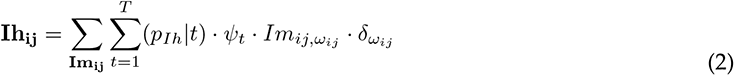

where, once again, *h* and *m* refer to hosts and mosquitoes and *I* and *S* refer to infected and susceptible individuals, respectively. For ease of interpretation we write this equation for one susceptible host species; multiple species could be represented with an additional subscript on each term. The outcome matrix **Ih_ij_** gives the total number of second generation hosts in each landscape cell [i, j] that become infected by the mosquitoes infected in Eq. 1. One mosquito-to-host transmission event is shown with the red arrow between the yellow mosquito and the solid human in Figure 1B. The total number of infected hosts in each cell [i, j] is given by two sums: a sum over the infected mosquitoes (**Im_ij_**) and a sum over these mosquitoes’ infectious period (*t*). Internal to these two sums are three terms that define mosquito-to-host transmission across the *ω_ij_* cells: (*p_Ih_|t*), the probability of mosquito-to-host transmission during blood feeding; *Im_ij,ωij_*, the number of infected mosquitoes from *Im_ij_* in *ω_ij_* ; and *δ_ωij_*, the biting rate of each infected mosquito on susceptible hosts of a given species. We now unpack these three terms.

Susceptible hosts in *ω_ij_* (we use *ω_ij_* to refer to the cell and *Ih_ij_* to the number of hosts that become infected in that cell) can become infected by being fed upon by infected mosquitoes. The mosquitoes from a given home cell (one entry of *Im_ij_*) will have the opportunity to feed on, and potentially infect, susceptible hosts in *ω_ij_* if their flight distribution includes *ω_ij_* (e.g., the yellow mosquito in Figure 1B includes the solid boxed cell with the human). The probability a mosquito infects a host during a feeding event increases over the mosquito’s infectious period because of the extrinsic incubation period of the virus within the mosquito; we write this as (*p_Ih_|t*), where *t* refers to the day post infection in the mosquito. The total number of feeding events (and thus potential host infections) by mosquitoes from *Ih_ij_* (one entry in the sum over **Ih_ij_**) on susceptible hosts in cell *ω_ij_* is first an increasing function of the number of infected mosquitoes (*Im_ij,ωij_*). The total number of feeding events by these mosquitoes on individuals of a given species of susceptible host (e.g., humans) is also a function of the feeding rate of each mosquito on that host species. This feeding rate is an increasing function of: the relative abundance of that species relative to other species, the mosquito’s intrinsic feeding preference on that species, and the mosquitoes’ general feeding rate. We collapse these three terms into *δ_ωij_* .

Our R_0_*-like* quantity is calculated for the cell in which the source infection originates (e.g., the red boxed cell in Figure 1A) as the sum of **Ih_ij_** across all host types that become infectious (which varies by disease, see details below). Thus, equations Eq. 1 and Eq. 2 show the complete calculation for R_0_ for one landscape cell. Conversely, the FOI for a specific landscape cell [i,j] (a cell defined by *ω_ij_* in Eq. 2) is calculated as the total number of hosts that become infected in that cell from infections starting in all landscape cells (for which Eq. 1 and Eq. 2 provide one example). Note that when calculated in this way, FOI is a breakdown of R_0_ values that are spatially reorganized and summed. Thus, over an entire landscape the sum of all R_0_ and all FOI values are identical.

We parameterized each component of Eq. 1 and Eq. 2 using estimates from either statistical models fitted to data extracted from published literature, or directly from parameter estimates in published literature; we describe all statistical models, data, and assumptions below. For all components of both transmission steps for all three diseases we searched the literature for raw data and quantitative parameter estimates (all references and raw data are available in the online supplement); when data were not available we relied upon qualitative, and occasionally anecdotal, descriptions (additional detail is available in the online supplement).

We considered the transmission of dengue by *Aedes aegypti* and *Ae. albopictus* and the transmission of malaria by *Nyssorhynchus darlingi*. An extensive literature search (details in the online supplement) made it clear that while many host species are fed upon by these mosquito species, and some of them are possibly able to become infected and transmit dengue and malaria, we know definitively very little about which non-human host species become infected with these pathogens and how competent they are in transmitting infection to susceptible mosquitoes. Further, the disease strains that non-human hosts are able to transmit are generally different than the strains that infect humans (Prugnolle et al., 2011, Maeno et al., 2015, Rondó n et al., 2019), which makes modeling the spillover of dengue and malaria a separate modeling task. Thus, for both of theses diseases we consider humans as the only host capable of transmission. However, because blood feeding by infected mosquitoes on non-human hosts will reduce disease burden on humans, it is important to consider non-humans in the model for these diseases. This is especially true for spatial estimates of disease risk as a function of landscape features because the proportion of an infectious mosquito’s bites on humans and non-humans will depend in part on the relative densities of these hosts (in addition to intrinsic mosquito preferences), which will vary across the landscape. For simplicity, computational efficiency, and data limitations we group all non-human species into a single type which we call “other” hosts.

We modeled both urban and sylvatic spillover of yellow fever by considering transmission by *Ae. aegypti*, *Ae. albopictus*, and *Haemagogus* spp.. To capture the predominant form of sylvatic transmission, which occurs between non-human primates and *Haemagogus* spp. (also *Sabethes* spp.) mosquitoes, we use humans as the primary host species, primates as a secondary host species, and “others” as hosts that mosquitoes feed upon but are unable to transmit the disease.

### Model components independent of landscape features

We parameterized most model components that we assumed to be independent of landscape features using estimates from statistical models fit to data extracted from the literature, including: host infection (titer) profiles over time (Eq. 1: *θ_dIh_*), mosquito infection probability (Eq. 1: *p_Im_*), mosquito transmission probability (Eq. 2: (*p_Ih_*), and mosquito survival (Eq. 2: *ψ_t_*). For host infection profiles, mosquito infection probability, and mosquito transmission probability we fit regression models to host and mosquito physiological responses to laboratory experimental infections. These models included fixed effects such as days since experimental exposure, infectious dose, and species. For host pathogen load we used a linear model and included a quadratic term for days since exposure to capture the rise and fall of pathogen load over a host’s infectious period; for mosquito infection and transmission probability we used generalized linear models with binomial error distributions. For all models we estimated responses (host pathogen load, mosquito infection and transmission probabilities) for host–pathogen and mosquito–pathogen pairs for which data were available, and relied upon qualitative descriptions in literature when quantitative data were unavailable. We show all fitted and assumed host infection profiles in Figure S1, mosquito infection probability curves in Figure S2, and transmission probability curves in Figure S3, and describe in the online supplement the data and assumptions used to generate each estimated response.

We used a simple exponential decay function to model the lifetime of infected mosquitoes (survival probability up to day X = *λ^X^*), where *λ* is daily survival probability. We gathered data from a small number of papers on the survival of each mosquito species (see Bates, 1947, Galindo, 1958, Dégallier et al., 1998, Muir and Kay, 1998, Niebylski and Craig Jr, 1994, Kiszewski et al., 2004, Maciel-De-Freitas et al., 2007, Lacroix et al., 2009, de Barros et al., 2011). Mosquito survival curves are shown in Figure S3; all raw extracted quantitative data for all mosquito species are available in the supplemental data files.

For mosquito feeding preference behavior (Eq. 1 and Eq. 2: part of the *δ* term) we simply used raw outcomes from blood meal analyses of wild-caught mosquitoes instead of fitting a model for mosquito biting preference. Specifically, given our approach of collapsing non-human host types, we collapsed the proportion of blood meals in wild mosquitoes into human and non-human sources. This approach does conflate mosquito species-specific intrinsic biting preference with the raw abundance of hosts, which could result in biased biting rates on our simulated landscapes; however, our simplified host model and lack of data made this simplification necessary. This simplifying assumption could be relaxed in a system with more species-specific mosquito biting behavior data available. For data sources and raw blood meal data see the online supplement.

### Model components dependent on landscape features

#### Host abundance and mosquito abundance

We assumed that human population density was directly proportional to urban intensity and that the density of non-human species (primates and “others”) were directly proportional to tree cover, with a scaling factor that can vary in order to adjust the absolute abundance of each host type on the landscape (additional detail available in the online supplement). While assuming different proportional relationships between urban intensity and human population density maybe somewhat unrealistic, we explore different proportional relationships so that we are able to isolate changes in human population density from landscape heterogeneity.

We modeled the relationship between the abundance of each mosquito species and both tree cover and human population density using details on the preferred breeding habitat of mosquitoes and where mosquitoes tend to be observed/collected in the wild (Braks et al., 2003, Scott and Morrison, 2010, Sarfraz et al., 2012, 2014, de Moura Rodrigues et al., 2015, Mucci et al., 2015, de Camargo-Neves et al., 2005, Lin et al., 2016, Tátila-Ferreira et al., 2017, Pereira dos Santos et al., 2018, Delatorre et al., 2019, Koyoc-Cardeñ a et al., 2019, Hendy et al., 2020, Silva et al., 2020). We assumed that each landscape cell supports a given resident population of each mosquito species (which could be thought of as their breeding location) based on the composition of that cell (in the case of *Ny. darlingi* also on the surrounding cells given their preference for forest edge: Tadei et al. 1998, Vittor et al. 2006, Zeilhofer et al. 2007, Vittor et al. 2009). We describe in detail the relationship between mosquito species abundance and individual landscape features in the online supplement and show mosquito densities on an example simulated landscape in Figure S4 (*Ae. aegypti*), Figure S5 (*Ae. albopictus*), Figure S6 (*Ny. darlingi*), and Figure S7 (*Haemagogus* spp.).

#### Mosquito movement and host movement

Our model for mosquito abundance assumes that each cell contains a resident population of mosquitoes of each species. It also assumes that each mosquito disperses from its home cell into the surrounding landscape cells during its lifetime for the purposes of blood feeding. We modeled the dispersal of mosquitoes from their home cells using a Gaussian spatial kernel with radius based on mark recapture experiments and observational field data (Causey and Kumm, 1948, Charlwood et al., 1995, Muir and Kay, 1998, Harrington et al., 2005, Russell et al., 2005, Achee et al., 2007, Vanwambeke et al., 2007, Hiwat and Bretas, 2011, Verdonschot and Besse-Lototskaya, 2014). We also included the option to weight this dispersal kernel by the habitat preference of each mosquito species (e.g., an *Aedes aegypti* mosquito with high human feeding preference that has a home cell on the border of a patch of urban area and a patch of high tree cover will spend more of its flight time in the urban area than under the high density tree canopy). However, we do not use this option in analyses presented here because: 1) we set the diameter of each landscape cell to equal the flight radius of an *Ae. aegypti*, which means that *Ae. aegypti* do not leave their cell; 2) we lack enough information on the remaining mosquitoes to accurately parameterize this weighting function. We also modeled the movement of the original infected host using a Gaussian kernel weighted in direct proportion to fraction urbanized.

### Landscapes

#### Simulated Landscapes

For the first stage of our analysis, we simulated landscapes composed of continuous values of tree cover and urban intensity on a scale of zero to one. To simulate varying degrees of landscape heterogeneity (measured with spatial auto-correlation: i.e., low landscape heterogeneity is simulated with high spatial auto-correlation) we used the ”midpoint displacement neutral landscape model“ (Barnsley et al., 1988), implemented in the package `NLMR` (Sciaini et al., 2018) (function `nlm mpd`) in `R` (R Core Team, 2020). With high spatial auto-correlation in each landscape feature, large areas of high tree cover are segregated from large areas of urban intensity (a land-sparing approach); with low spatial-auto-correlation features are highly spatially mixed (a land-sharing approach). For each desired level of landscape heterogeneity we generated landscapes with ecologically realistic patterns of tree cover and urban intensity using the following procedure: 1) simulate two individual landscape matrices with values ranging from zero to one, one matrix of tree cover and one matrix of urban area; 2) check if the average value for each matrix falls outside of *x ± E_x_*, where we set *x* = 0.50 (a balanced landscape) and *E_x_* = 0.02 (as narrow as possible while maintaining computational efficiency); 3) check if the correlation between the values in the two matrices falls outside of *y ± E_y_*, where we set *y* = *−*0.50 and *E_y_* = 0.02 (with this range, the highest values of forest cover will tend to not occur in highly urbanized cells, though urban areas and tree cover will overlap to a moderate degree); 4) if any of the values fall outside of the desired ranges, repeat from step one. Four example simulated landscapes across the full range of auto-correlations we used are pictured in Figure S8.

Although we did not directly simulate additional types of landscape features (such as agriculture), areas of our simulated landscape with minimal urban intensity and minimal tree cover can be interpreted as an agriculture-like habitat. We took this approach for two reasons. First, little quantitative information exists on the relationship between the abundance of *Ae. aegypti*, *Ae. albipictus*, and *Ny. darlingi* and other landscape features that could be abstracted in a simulation but still be used to parameterize mosquito abundance. Second, simulating three matrices with appropriate levels of spatial auto-correlation, negative correlation with each other, and average values would be difficult.

#### Empirical Landscapes

For the second stage of our analysis we selected a small landscape (approximately 400 km^2^) just to the northwest of Bogotá, Colombia, which is part of a larger area of Latin America that is of high restoration importance (Strassburg et al., 2020) (Figure S9). This region contains a portion of a major city, intensive agriculture, towns, rural homesteads, and forest. For this landscape we extracted human population density as of 2010 (Sorichetta et al., 2015, WorldPop, 2016), and leaf area index (LAI) in 2013 as a proxy for tree cover (Gerard et al., 2020).

We simulated habitat restoration on this landscape, modeled as an increase in LAI in three different ways: 1) reforestation of a single large contiguous area (simulated with `NLMR` using high spatial auto-correlation); 2) patchy reforestation across the landscape (e.g., reforestation on individual farms; simulated with low spatial auto-correlation); and 3) “flat” reforestation which we modeled as an increase in the LAI in all cells in proportion to their sampled values. For both restoration scenarios one and two, we simulated reforestation under the constraint that the post-restoration simulated landscapes had a minimum negative correlation between human population density and tree cover of -0.45 (as we did with the simulated landscapes in stage one of our analysis). For each scenario we modeled an increase in average LAI on the whole landscape from baseline (0.17) to a value of 0.50, the same average used in our stage one simulated landscapes.

To predict how disease transmission would change as a function of restoration strategy, we compared estimated FOI on the baseline landscape to FOI estimated for each restoration scenario. This simulation ignores the complex ecological process (and time lag) of colonization of the newly restored forest by hosts and mosquitoes, and simply assumes that the newly restored forest contains “other” hosts, non-human primates, and forest associated mosquitoes in proportion to the assumed relationship between tree cover and host and mosquito abundance used to predict baseline disease transmission.

## Results

### Simulated Landscapes

On simulated landscapes of urban area and tree cover, average human infection risk (FOI_h_) for dengue and yellow fever is driven primarily by human population density (Figure 2, with effects in differing directions), while malaria risk is a function of both human population density and the spatial configuration of urban area and tree cover (Figure 2). Here, and in the rest of the results we focus on spatial patterns and summaries of FOI_h_ instead of R_0_ (or FOI on non-human primates or ”others“) to focus on those landscape regions for which the potential disease flow into humans is highest (recall that FOI is just a spatial reorganization of R_0_ values and that across an entire landscape total FOI is equal to total R_0_ as both metrics rely on the assumption of a source infection appearing in each landscape cell).

**Figure 2:**
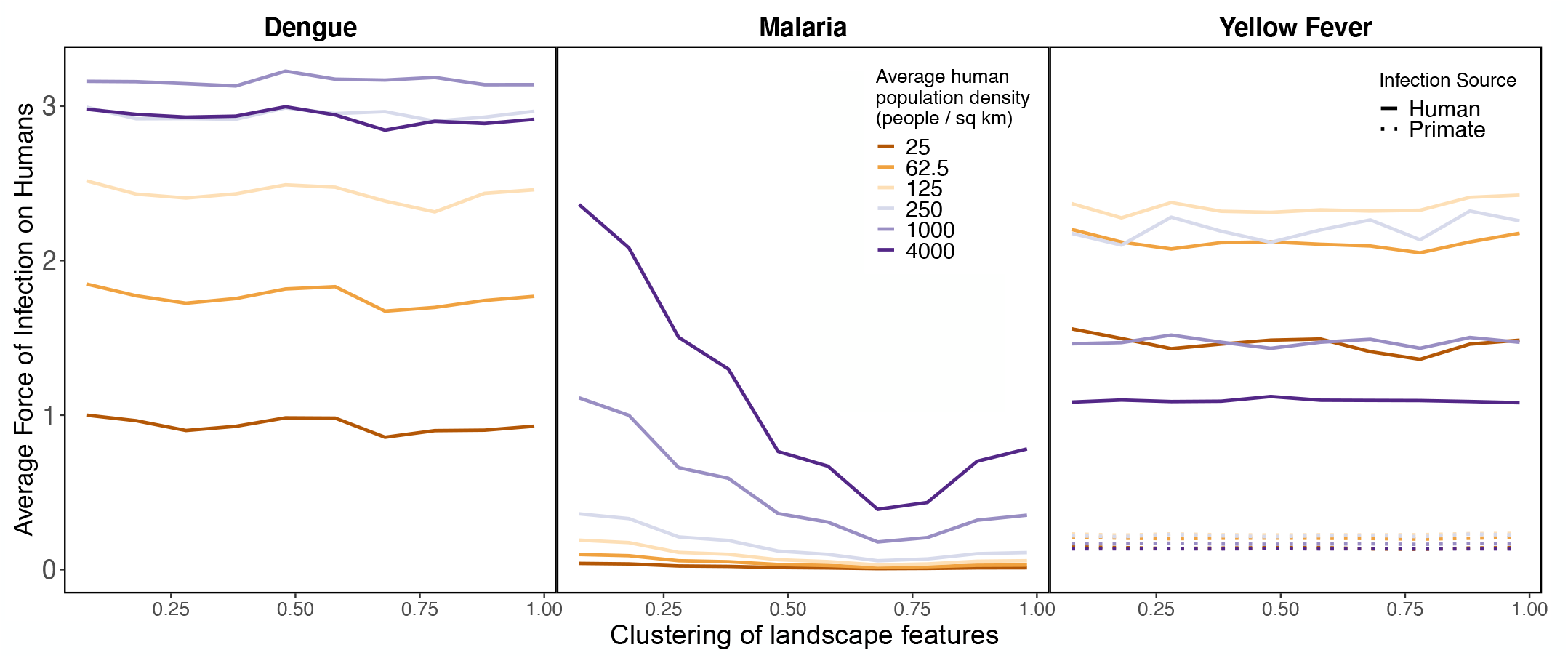
Human density drives variation in the average force of infection on humans for dengue, malaria, and yellow fever; landscape configuration affects malaria FOI_h_ nonlinearly. Colors show average human population density on the landscape. The clustering of landscape features (x-axis) refers to the values of the spatial auto-correlation used in simulating landscape-features; more clustering equates to decreasing landscape spatial heterogeneity (larger contiguous areas of high urban intensity and tree cover). The FOI_h_ shown here is an average of the FOI_h_ on all landscape cells, each of which is calculated by summing the FOI_h_ from source infections appearing in each landscape cell.

#### Dengue

Average dengue FOI_h_ across a landscape is a non-monotonic function of the human population density on that landscape (Figure 2). This pattern arises because of the combined transmission from *Ae. aegypti*, whose abundance is strongly tied to human population density and who prefer to feed on humans, and *Ae. albopictus*, whose abundance we assumed to be tied to tree cover but who also prefer to bite humans. As human population density increases, the abundance of *Ae. aegypti* increases, leading to a larger FOI_h_ attributable to *Ae. aegypti* (Figure 3). At the same time, however, because *Ae. albopictus* abundance is assumed to be independent of human population density, the single source human dengue infection becomes “lost” in a sea of susceptible humans to the constant population of blood feeding *Ae. albopictus*, leading to fewer infected *Ae. albopictus* (Figure S10) and subsequently fewer second-generation human infections (Figure 3). However, at very low human population densities the FOI_h_ from *Ae. albopictus* increases because of a higher proportion of infectious bites on humans relative to “other” species. Considering transmission by both *Ae. aegypti* and *Ae. albopictus*, average dengue infection risk is maximized at an intermediate human population density (Figure 3), though FOI_h_ decreases very minimally after the maximum. This highlights that, depending on the relative abundance and importance of *Ae. aegypti* versus *Ae. albopictus* in a given landscape, different levels of human density and forest cover could maximize dengue transmission.

**Figure 3:**
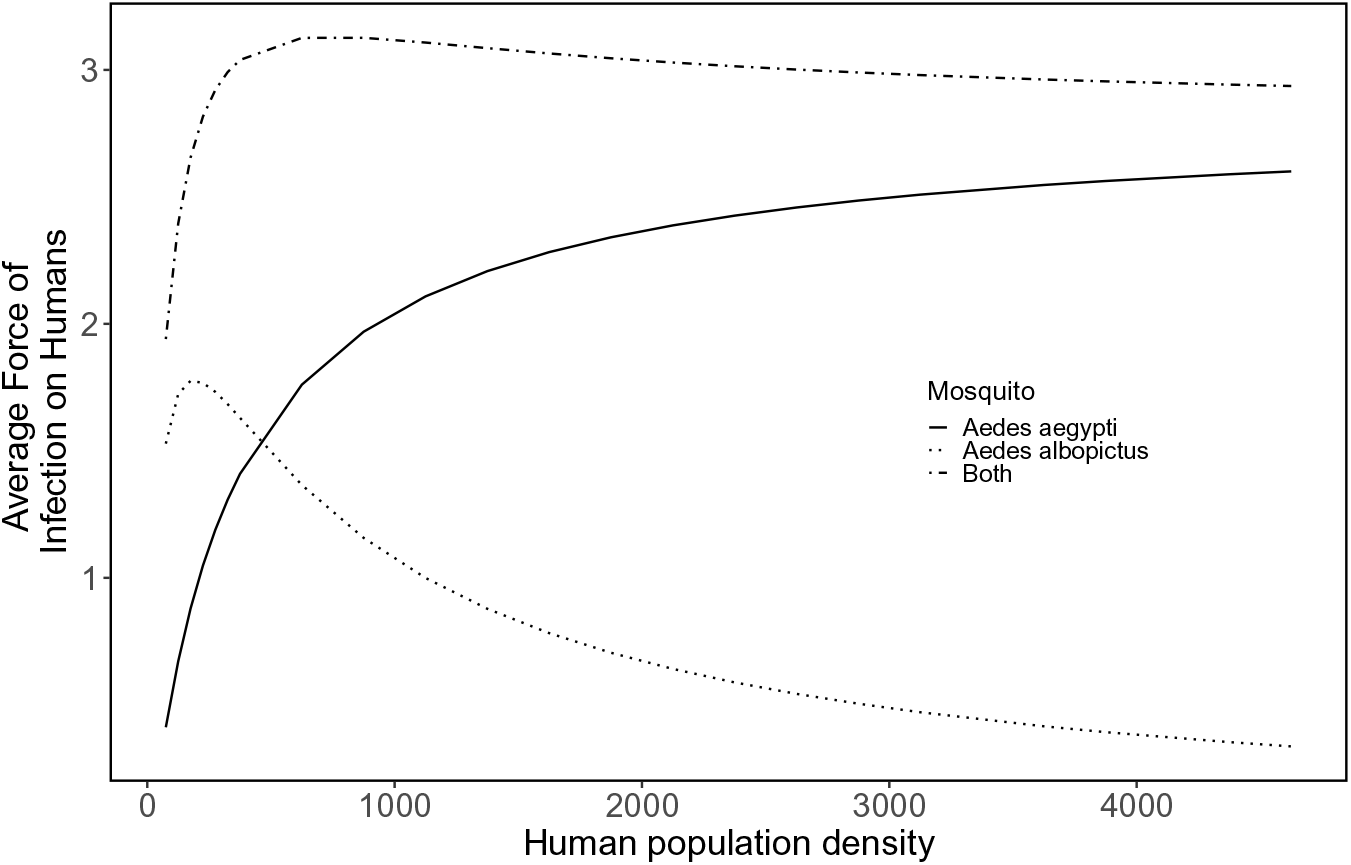
***Aedes aegypti* and *Aedes albopictus* combine to drive dengue FOI_h_ on simulated landscapes that vary in absolute human population density.** Total FOI_h_ is the sum of the contributions made by *Aedes aegypti*, which increases monotonically with human density, and *Aedes albopictus*, which peaks at low human density. These results were calculated on a landscape with a spatial auto-correlation of 0.78.

The relative contribution *Ae. aegypti* and *Ae. albopictus* make to overall human dengue risk also depends strongly on the relationship between *Ae. aegypti* abundance and human abundance. In Figure 2 and Figure 3 we assumed a linear relationship between human abundance and *Ae. aegypti* abundance; however, it has been suggested that this relationship may be exponential (Romeo-Aznar et al., 2018), such that the ratio of *Ae. aegypti* per human increases as human population size increases. Our simulations show that as this exponent increases from below one (a decreasing mosquito-to-human ratio with an increasing human population density) to above one (an increasing mosquito-to-human ratio), the contribution that *Ae. aegypti* make to FOI_h_ increases (Figure 4). At low human population densities the importance of *Ae. albopictus* is greater than the importance of *Ae. aegypti* (Figure 3, Figure 4); however, as human population density and the exponent linking human and *Ae. aegypti* populations increase, the importance of *Ae. aegypti* overtakes that of *Ae. albopictus*.

**Figure 4:**
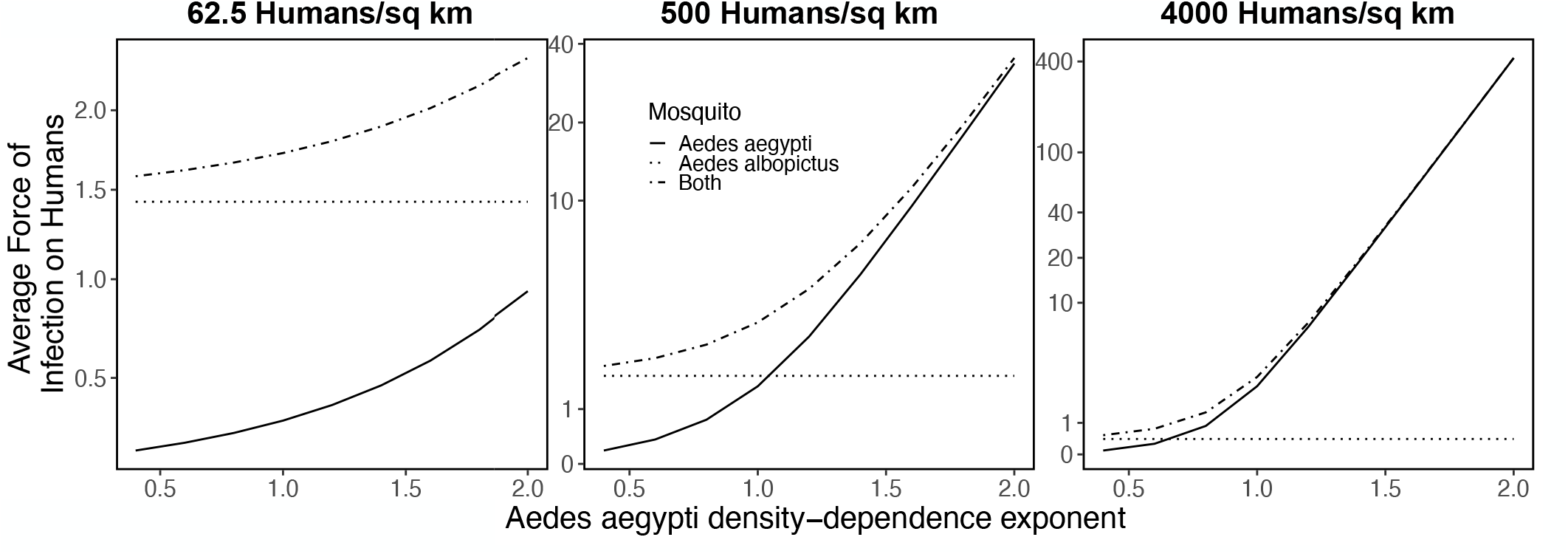
The contribution *Aedes aegypti* and *Aedes albopictus* make to dengue FOI_h_ as a function of human density and the exponential relationship between human abundance and *Aedes aegypti* abundance. Panels show low (62.5), medium (500), and high (4000) human population density (humans/sq km). Total FOI_h_ is the sum of the contributions made by *Aedes aegypti* (solid line) and *Aedes albopictus* (dotted line). Here we only manipulated the exponential relationship between human abundance and *Aedes aegypti*, thus the FOI_h_ attributed to *Aedes albopictus* remains constant. These results were calculated on a landscape with a spatial auto-correlation of 0.78.

#### Yellow Fever

Similar to dengue, yellow fever FOI_h_ is a non-monotonic function of human population density assuming either a source infection in a human (Figure 2, solid lines) or a source infection in a non-human primate (Figure 2, dotted lines). As with dengue, these patterns are due to the combined transmission by all mosquitoes involved; for yellow fever these mosquitoes include *Ae. aegypti*, *Ae. albopictus*, and *Haemagogus* spp. However, unlike for dengue (where *Ae. aegypti* begins to dominate transmission at relatively low population densities: Figure 3), an overall higher relative contribution by *Ae. albopictus* to transmission paired with an overall declining importance with increasing human population density (except at the very lowest human population densities) makes yellow fever FOI_h_ peak at a low human population density (Figure S11). Assuming a human source infection, yellow fever FOI_h_ is higher in regions of higher forest cover because it is transmitted by two forest-dwelling mosquitoes, while dengue increases with increasing urban intensity (Figure S12).

The number of spillover yellow fever infections from a source primate infection into humans is smaller than from a human infection (Figure 2), as spillover requires an initial blood feeding event by a mosquito on the infected primate followed by a blood feeding event on a susceptible human, which is rare for mosquitoes that feed preferentially on either humans (*Ae. aegypti* and *Ae. albopictus*) or non-humans (*Haemagogus* spp.). Similar to human-to-human yellow fever transmission, our model estimates that spillover FOI_h_ increases with decreasing human population density (though the effect size is quite small: Figure 2). Unlike for human-to-human transmission, increasing spillover transmission with decreasing human population density is driven by a higher probability of an initial feeding event by *Ae. aegypti* and *Ae. albopictus* on the infected primate when humans are rare. The effect size is small because most transmission from primates to humans is driven by *Haemagogus* spp., for which this relationship does not hold. Spillover transmission is mostly independent of landscape heterogeneity because spillover tends to occurs within landscape cells with low to moderate tree cover and population density, which is approximately constant among the simulated landscapes. Although analyses based on R_0_ (including our FOI_h_ calculation) that do not consider the impact of vaccination predict that human-originating infections have a larger FOI_h_, in populations with high yellow fever vaccination rates, spillover infections are likely to be the more important driver of infections in humans.

#### Malaria

Across all simulated landscapes we find a non-linear (and non-monotonic) relationship between landscape heterogeneity and average malaria FOI_h_, such that average malaria FOI_h_ is minimized at a high but sub-maximum spatial auto-correlation (*∼*0.70, see Figure 2). On individual landscapes we also find non-monotonic relationships between tree cover and malaria FOI_h_, and the shape of this relationship depends on the degree of landscape feature spatial auto-correlation (Figure 5D). On all landscapes, malaria FOI_h_ is maximized in areas of higher urban intensity within or adjacent to a region of patchy tree cover (as *Ny. darlingi* abundance is driven strongly by forest edge: *Methods: Mosquito abundance*, Figure S6). However, the composition and spatial structure of this interface changes with landscape feature spatial auto-correlation. On landscapes with moderate or high spatial auto-correlation (moderate or large contiguous patches of tree cover and urban area), high tree cover occurs in large patches that are distant from urban areas, while low tree cover occurs in large patches of urban area; in both cases FOI_h_ is low (Figure 5D). It is within the transition zone between dense tree cover and low tree cover that “forest edge” near high urban intensity exists, and thus where FOI_h_ is maximized. In contrast, on landscapes with low spatial auto-correlation (highly spatially integrated landscapes), which are characterized by small patchy urban areas and tree cover, small regions (even single cells) of high urban intensity can occur directly next to small patches of forest cover. First, this leads to an overall higher average malaria FOI_h_ across the landscape (Figure 5C) because of the larger number of humans within the flight radius of suitable *Ny. darlingi* habitat. Second, it causes FOI_h_ to be maximized in landscape cells with low tree cover and high urban intensity (Figure 5D). On highly heterogeneous landscapes (Figure 5, left column) these cells are commonly found within a broader region of mixed tree cover, which increases *Ny. darlingi* abundance in the area. Within the local region of suitable *Ny. darlingi* habitat, human infections will be concentrated in cells with low tree cover and high urban intensity as infectious bites on humans by dispersing *Ny. darlingi* will be the highest where the ratio of humans to other potential blood meal sources is maximized. These cells can be seen as red pixels in Figure 5 (Panel C, left column) and the small cluster of data points in the top left of Figure 5 (Panel D, left column). Finally, given that a higher human population density leads to a higher proportion of infectious bites on humans on any landscape, average malaria FOI_h_ (starting with a source infection in a human) is a monotonically increasing function of human population density (Figure 2, Figure 5D).

**Figure 5:**
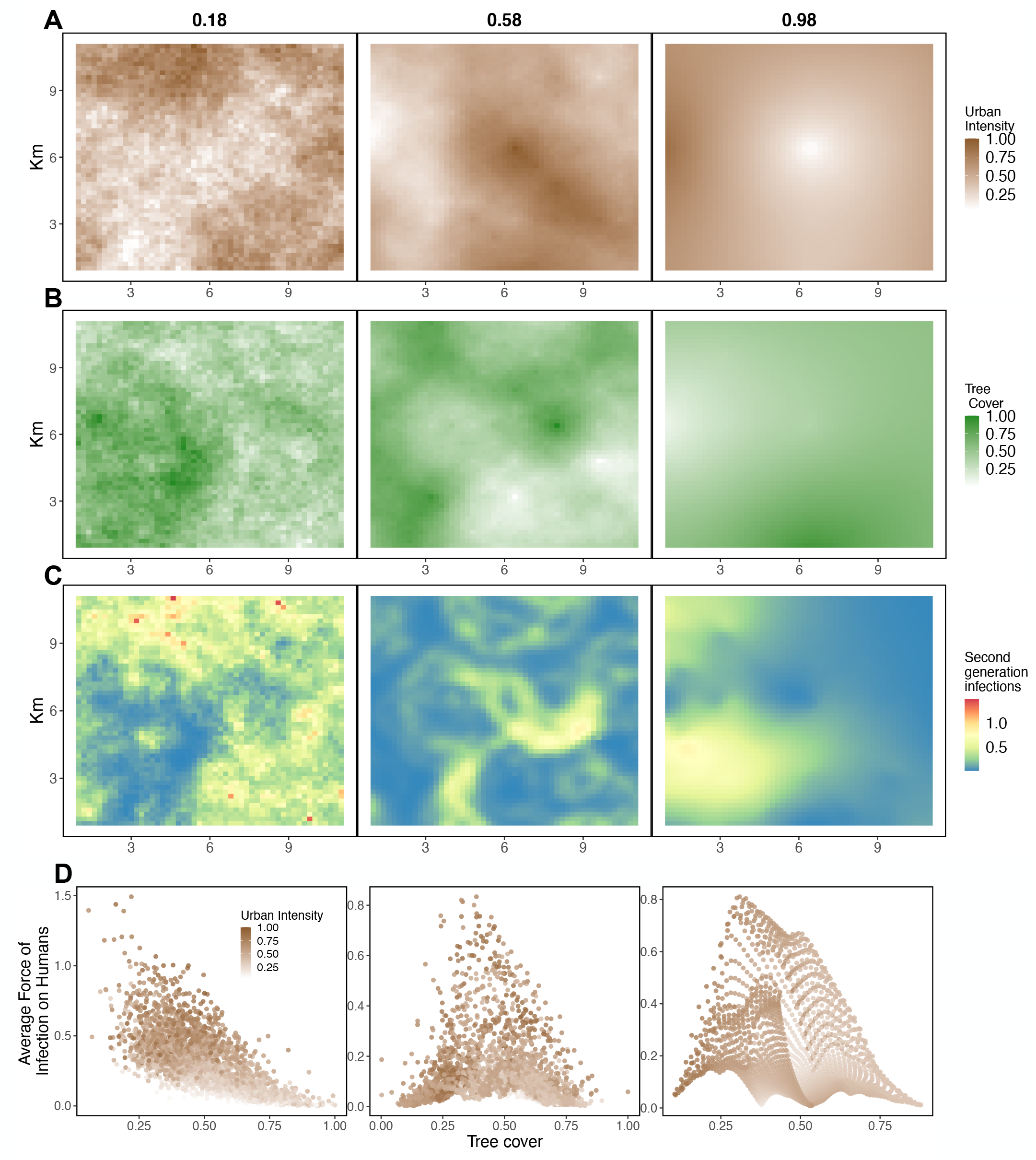
**Malaria FOI_h_ (Panel C) on simulated landscapes with low (0.18) medium (0.58) or high (0.98) spatial autocorrelation for urban area (Panel A) and tree cover (Panel B)**. The relationship between urban intensity (point color), tree cover (x-axis), and malaria FOI_h_ (y-axis) for all landscape cells across the three simulated landscapes are shown in Panel D. The relationship between tree cover and malaria FOI_h_ changes with the spatial autocorrelation, from monotonically negative in very patchy (low autocorrelation) landscapes to unimodal in moderate to high autocorrelation landscapes. The landscapes pictured here have an average population density of 250 people per sq. km.

#### Tradeoffs and synergies among diseases

Considered together, the FOI_h_ of dengue is positively correlated with malaria FOI_h_ across landscapes, though the strength of this correlation depends on the degree of spatial auto-correlation of landscape features and on the amount of forest cover and urban intensity (Figure 6). On highly heterogeneous landscapes (low spatial auto-correlation in Figure 6), dengue and malaria FOI_h_ are strongly positively correlated, though this correlation decreases with decreasing heterogeneity (Figure 6). The correlation tends to be slightly weaker in regions of the landscape with lower tree cover. Alternatively, the FOI_h_ of yellow fever and malaria is always moderately negatively correlated; decreasing heterogeneity reduces the size of this negative correlation only marginally, and subsetting to landscape regions with high or low urban intensity or tree cover has little effect (Figure 6). Finally, the correlation in FOI_h_ between dengue and yellow fever is often negative, but can be positive on landscapes of high spatial heterogeneity (low landscape feature spatial auto-correlation) in landscape cells with higher urban intensity and lower tree cover (Figure 6, Figure S12). On highly heterogeneous landscapes, individual cells with high urban intensity and low tree cover are commonly found within a broader region of higher tree cover; these cells experience higher dengue FOI_h_ because of larger *Ae. aegypi* populations and more yellow fever because of the dispersing *Ae. albopictus* and *Haemagogus* spp. from the surrounding area.

**Figure 6:**
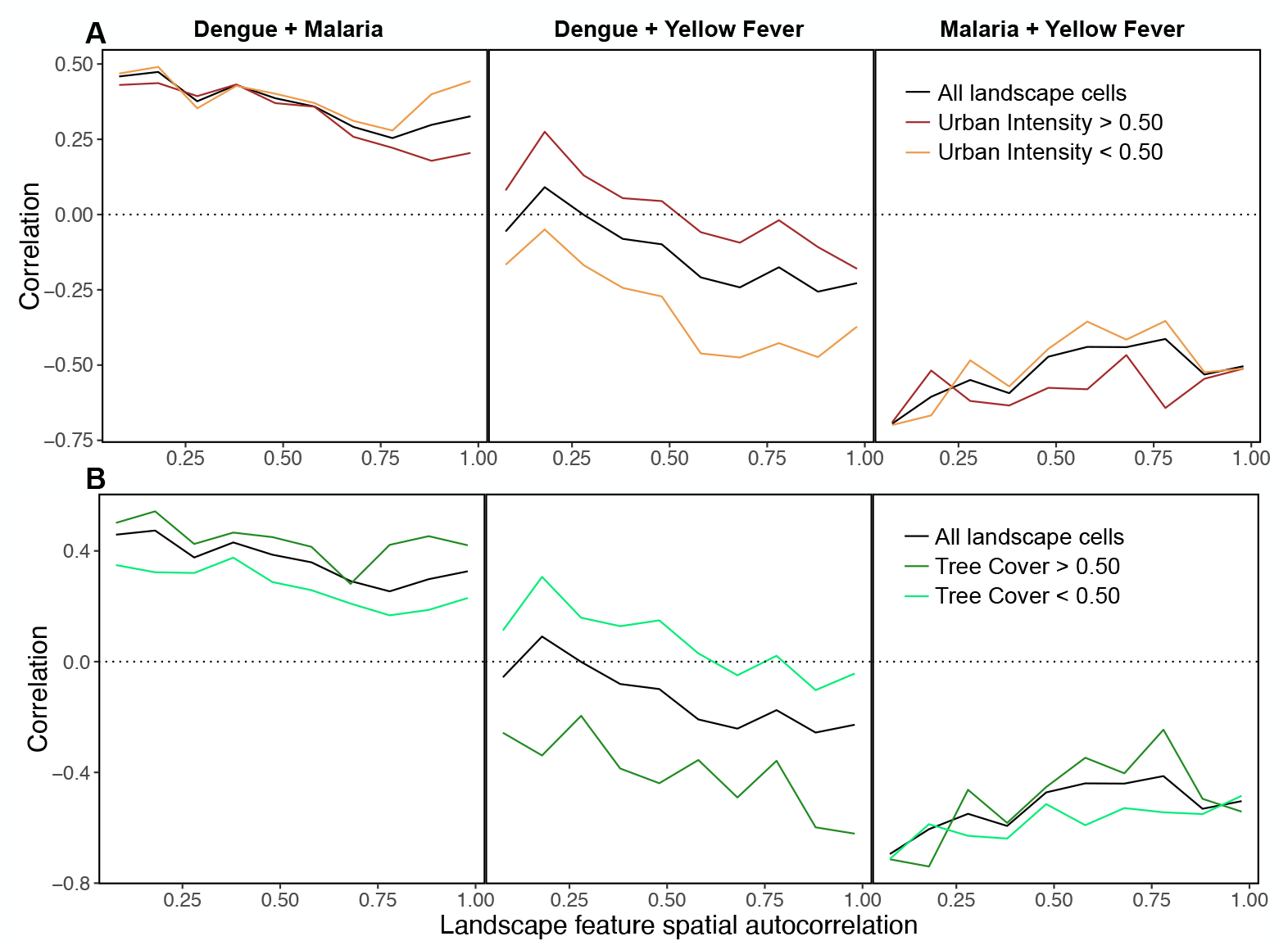
Malaria tends to be positively correlated with dengue and negatively correlated with yellow fever, while the correlation between dengue and yellow fever varies across urban and forested landscapes. Black lines in all panels show correlations across all cells on the landscape, while the darker brown and green lines in panels A and B show correlations between each disease in landscape cells with greater than 0.50 urban intensity and tree cover, respectively. Similarly, lighter colored brown and green lines in panels A and B show correlations between each disease in landscape cells with less than 0.50 urban intensity and tree cover, respectively. All results pictured here are for landscapes with average human population density of 250 people per sq.km. (Correlations are identical across densities; not pictured.)

While it is more difficult to parse the quantitative relationships among FOI_h_ when viewed in the form of map layers, stacked map layers are useful to illustrate that each disease has its own spatial pattern and that single local regions of a heterogeneous landscape are unlikely to have a high FOI_h_ for all diseases (Figure 7). For landscapes with either moderate (e.g., Figure 7), low, or high spatial heterogeneity, total disease risk (sum of FOI_h_ values for each disease) is maximized at intermediate to high values of urban intensity and tree cover because of positive correlations between both dengue and yellow fever FOI_h_ and urban intensity and tree cover (Figure S13). Though malaria is maximized at intermediate values of both landscape features (Figure S13), malaria has an overall smaller estimated FOIh and thus contributes less to total FOIh (Figure S14).

**Figure 7:**
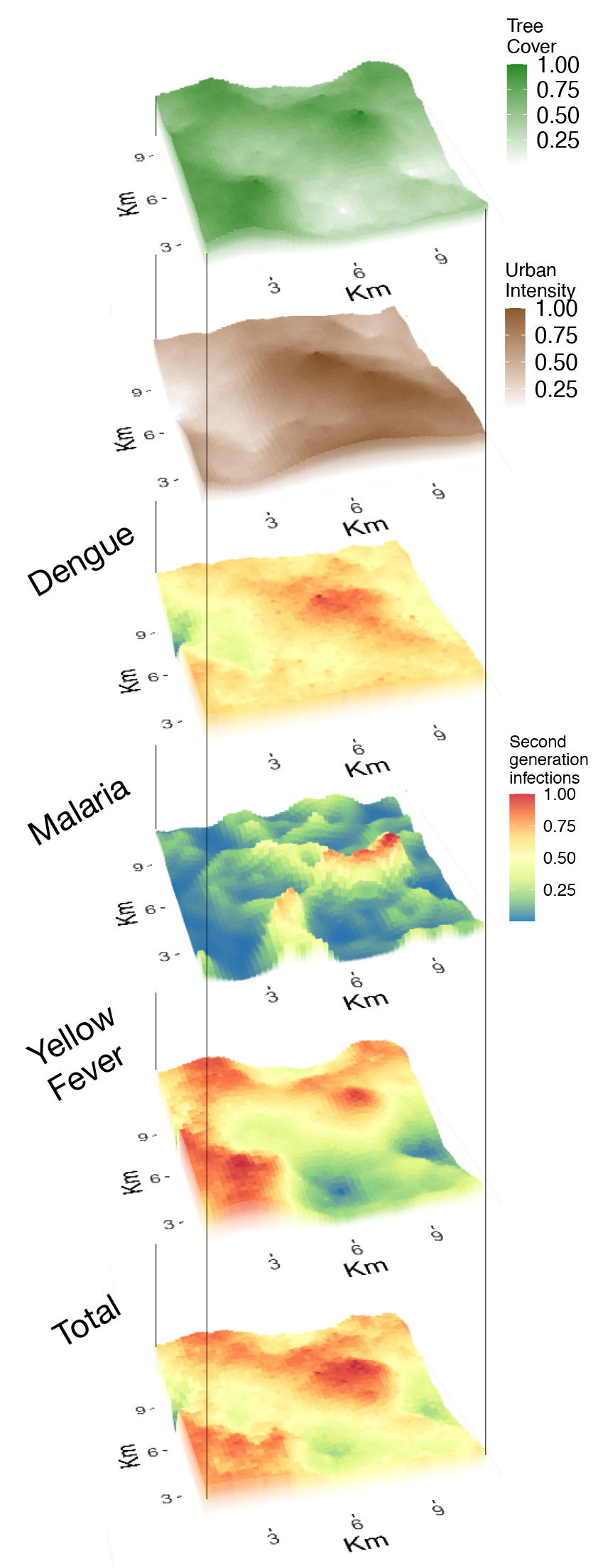
Simulated landscape features and estimated FOI_h_ of each disease, aligned and stacked to aid visualization of the overlap of features with high-risk and low-risk regions of the landscape. FOI_h_ values for each disease are on a relative scale (0, 1) to focus on the spatial patterns in disease risk (absolute FOI_h_ values are shown in Figure S14). This simulated landscape is the same as that with medium spatial autocorrelation among landscape features (0.58) shown in Figure 5, which has an average human population density of 250 people per sq.km.

### Reforestation Scenarios on a Real Landscape

While estimates of FOI_h_ on simulated landscapes can help to reveal general patterns in disease risk, it is unclear how well such patterns translate to real landscapes. To connect our model more strongly to a real-world scenario, we estimated the FOI_h_ of each disease on a 23km x 18km landscape to the northwest of Bogotá, Colombia that is a heterogeneous mixture of urban area and farmland, and has some, but an overall low average tree cover (average LAI across the landscape of 0.18). From this baseline we calculated disease risk as a function of three reforestation scenarios: “Flat” increases tree cover evenly across the landscape (which serves as a null model), “Congtiguous” simulates the planting of a single large patch of forest (e.g., a regional conservation effort), and “Patchy” simulates the planting of many small patches (e.g., subsidies to individual farms to replant trees).

For both dengue and yellow fever we predict a small increase in FOI_h_ under all reforestation scenarios (Figure 8), though we predict the largest increase in risk under patchy reforestation because of increased contacts between humans and *Ae. albopictus* and *Haemagogus* spp. All reforestation scenarios have a relatively small impact on average malaria FOI_h_ across the whole landscape (because of large regions of low FOI_h_: Figure 8, Panel C, blue regions), though the “Flat” and “Patchy” scenarios do introduce a series of new risky host-spots (Figure 8C). For example, under a “Flat” scenario, these areas are concentrated in the southeastern and western part of the landscape, which are areas that at baseline are urban edges near sparse forests.

**Figure 8:**
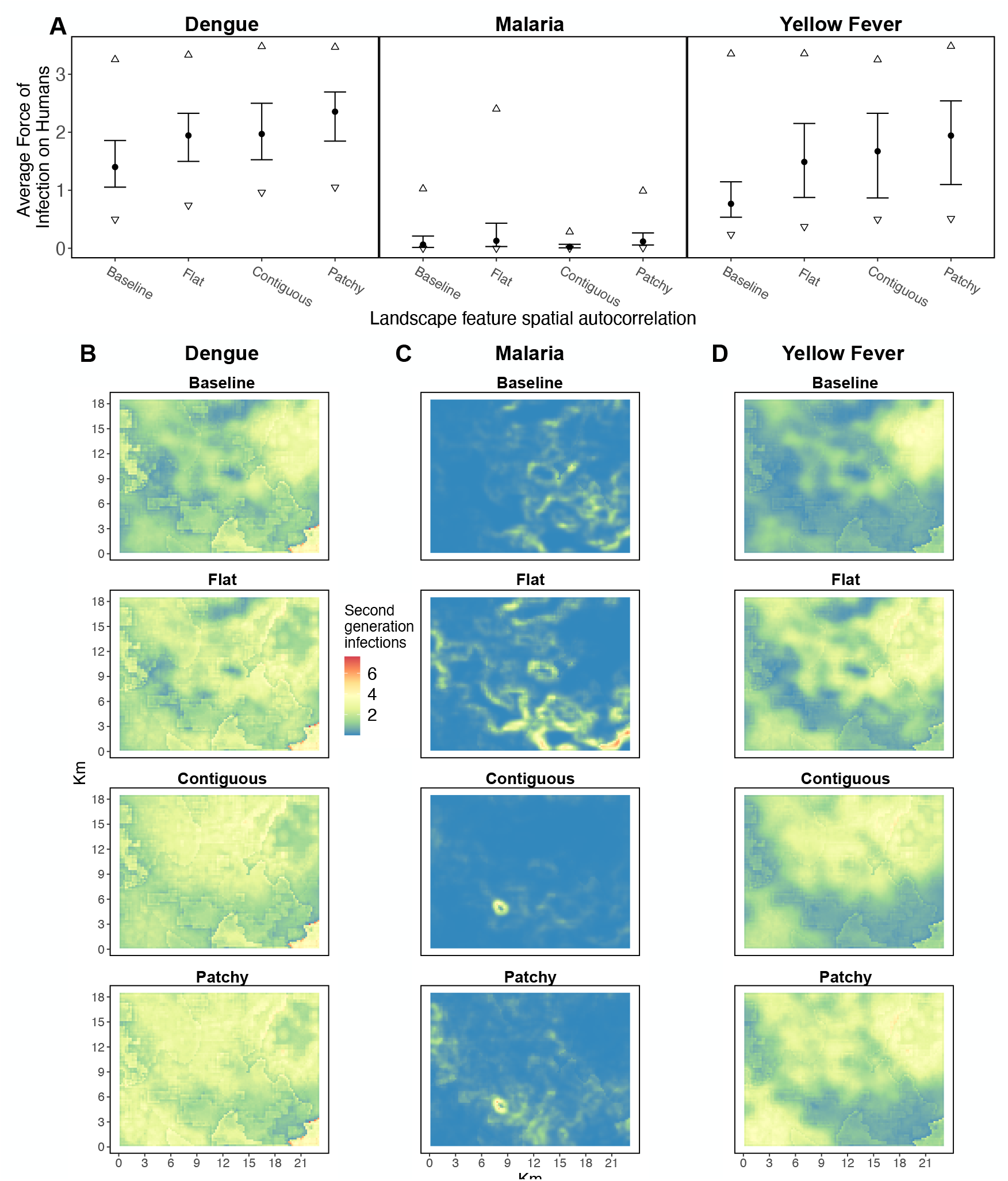
**Estimated FOI_h_ of each disease on a 23km x 18km landscape to the northwest of Bogotá, Colombia for three potential scenarios of reforestation**. Panel A shows the estimated average FOI_h_ of each disease (error bars show the central 50% of values while triangles show central 95% of values) on the landscape as it appears (“Baseline”) (see the online supplement for details about the data and Figure S9 for a map of the region) and for three reforestation scenarios: “Flat” increased tree cover evenly across the landscape (which serves as a null-model), “Congtiguous” simulated the planting of a single large patch of forest (e.g., a regional conservation effort), and “Patchy” simulated the planting of many small patches (e.g., subsidies to individual farms to replant trees). For all scenarios the average tree cover on the landscape is simulated to be brought up from 0.14 (as measured by LAI; see supplemental methods) at baseline to 0.50. Panels B-D show maps of the estimated FOI_h_ of dengue, malaria, and yellow fever, respectively, for baseline and each reforestation scenario; the shape in bottom right faintly outlined in red in panel B is the northwest corner of Bogotá (see Figure S9).

As an alternative to the somewhat contrived assumption of infections arising on each landscape cell, FOI_h_ can alternatively be modeled assuming that only a *single* infection of each disease were to appear somewhere on the landscape (where infection emergence is weighted by, for example, human population density). Assuming this alternative definition for FOI_h_ produced qualitatively similar results (Figure S15) to those presented here.

## Discussion

With high rates of land-use change globally (Seto et al., 2013, Runyan and D’Odorico, 2016, Dinerstein et al., 2019, UN, 2019) and mounting evidence that land-use change affects the transmission of infectious diseases (Sharma, 2002, Ward and Brown, 2004, Vanwambeke et al., 2007, Hahn et al., 2014, Sheela et al., 2017, Ziemann et al., 2018, MacDonald and Mordecai, 2019), it is important to seek a stronger mechanistic understanding of the link between land-use patterns and disease risk. The ability to predict how future land-use change will affect human health would help to inform interventions (e.g., where to apply mosquito control) and to design restoration strategies that minimize risk. Here, we designed and analyzed a spatially explicit model of disease transmission to predict how the spatial configuration and density of tree cover alongside urban area and human population density affects the potential for disease transmission (using R_0_) and where human risk of disease (FOI_h_) is highest.

This model was intended, first and foremost, to provide a road map for how the multifaceted impacts of land-use on disease transmission can be mechanistically modeled to understand and predict changes in disease risk in response to land-use change, including both degradation and restoration. Our analysis sought to conceptualize how different types of diseases with different transmission strategies—parameterized to represent dengue, malaria, and yellow fever—would respond to different landscape configurations and land-use change scenarios. At the broadest level, our results serve as a valuable proof of concept that different, human-important diseases depend on the biophysical landscape in different nonlinear ways (Figure 2, Figure 5, Figure S13), leading to correlations among diseases that are themselves not constant across the landscape (Figure 6, Figure 7). This complexity and nuance suggests that relying on simple “rules of thumb” for the relationship between land-use and disease could lead to sub-optimal or even dangerous planning/restoration decisions; a model that incorporates these nonlinear mechanisms, such as the one presented here, will be a prerequisite for applied research on ecosystem services for health. We found, for example, that both dengue and yellow fever were highly dependent on human population abundance and less so on the spatial configuration of the landscape, though dengue FOI_h_ increases with urban intensity while yellow fever FOI_h_ is maximized in areas of moderate urban intensity and high tree cover (Figure S13). In contrast, we estimated that malaria was highly dependent on the spatial configuration of urban area and tree cover, and that human risk of malaria peaked at the interface of urban areas and forest when there was a high variance in tree cover (forest edge and not large, high density swaths of forest).

While we found that different forest restoration strategies can have different impacts on disease risk in a spatially heterogeneous way, we did also find similarities in how patchy vs contiguous reforestation impacted the landscape-level average risk for each disease (Figure 8). For example, we estimated that patchy reforestation, akin to a “land-sharing” strategy, on a small landscape to the northwest of Bogotá, Colombia could increase the risk of all three diseases on average, though much of the increased risk would be borne by individuals living in small urban areas adjacent to the increased tree cover who would be expected to experience higher rates of infection from *Ae. albopictus* and *Ny. darlingi*. Alternatively, we showed that a single large contiguous patch of increased tree cover, akin to a “land-sparing” strategy, would decrease average malaria risk, reduce the number of high risk malaria hotspots (Figure 8), and lead to a smaller increase in dengue and yellow fever risk. While we have strong empirical evidence for increased forest fragmentation leading to increased malaria (Vittor et al., 2006, Hahn et al., 2014, MacDonald and Mordecai, 2019), our results suggest it will also be important to monitor the reverse of this trend.

As with any model, a number of our underlying assumptions influenced model predictions: in particular, the structure of the functional forms we assumed and the parameter values we used for those functional forms. Though we conducted an extensive literature search for each model parameter, we often failed to find quantitative estimates that would allow us to parameterize relationships between urban intensity or tree cover and transmission-related quantities like mosquito abundance. For example, in the absence of direct empirical data, we translated a qualitative understanding that *Ny. darlingi* prefer forest edge habitat into a quantitative link between spatial heterogeneity in tree cover and *Ny. darlingi* abundance. Even for *Ae. aegypti*, which is extensively studied because of its importance in transmitting dengue, Zika, chikungunya, and yellow fever, we still know little about the quantitative relationship between its abundance and human abundance (Romeo-Aznar et al., 2018). Given the strong dependence of dengue on the relationship between human abundance and *Ae. aegypti* abundance we find here (Figure 4), this is a priority area for future empirical work. Further, given a lack of sufficiently detailed mosquito blood meal data, we used a simplified representation of mosquito feeding behavior. Because mosquito feeding behavior affects both host-to-mosquito transmission and mosquito-to-host transmission, it has a large impact on results; further empirical work on the feeding preferences of these mosquitoes would help to improve model estimates.

Given the uncertainty in functional forms and parameter values governing relationships between land use and disease, these assumptions should be refined within a local context before applying this approach directly to decision-making. In the meantime, we suggest that the model could be used as a tool for analyzing uncertain phenomena and forming hypotheses for future testing (Baker et al., 2018). For example, we have shown that the model can provide an early expectation for the broad range of effect sizes that various reforestation strategies could have (Figure 8). It can also be used to examine more nuanced patterns, such as the prediction that in the presence of both an urban-breeding mosquito and a forest-breeding mosquito that both prefer biting a host that dwells primarily in one landscape area (which are represented here by *Ae. aegypti*, *Ae. albopictus*, and humans), the FOI_h_ attributable to each mosquito will be inverses of one another leading to a peak in infection risk at intermediate host population density. However, it may also be possible to circumvent these data limitations if a time series of spatially explicit human disease incidence data is available (e.g., from health centers spread across the focal landscape). With these data, unknown model parameters could be calibrated by matching predictions to the spatial health records. Optimally, calibration would happen over time by running the model from some time in the past until the present using a time series of land-use snapshots (e.g., using remote sensing derived LAI). Following this calibration, the model could then be run into the future with various scenarios of potential land-use change.

To realize this use, however, two roadblocks would need to be overcome. First, any parameter calibration would require human disease notifications recorded spatially over time, such as from a detailed health surveillance system. Second, even with these data in hand, a modification may have to be made to link reported cases to R_0_ and FOI. Further, a few additional model caveats will be important to keep in mind. First, we assumed a simple Gaussian spatial pattern of movement of the infected host around a “home” landscape cell. If movement of the infected host or vector is more complicated (such as an infected human moving long distances along a road, between a few specific focal points of interest, or alternatively a sick individual not moving at all; e.g., see Stoddard et al. 2013, Kennedy et al. 2016), spatial FOI patterns could look very different (e.g., much flatter across the landscape if movement is much wider, or more patchy if movement is lower). We also assumed that the infected individual’s availability to mosquito feeding is constant over their infectious period, which is a simplified version of infection dynamics. For example, humans can transmit dengue to people inside or outside their households, and a varying proportion of transmission occurs before versus after symptom onset depending on how illness modifies behavior (Sch- aber et al., 2021). Finally, because the model is designed to calculate snapshots of risk on static landscapes, it assumes instantaneous ecological succession; that is, all that determines host and mosquito abundance on the landscape is the features of the current landscape and not the past history of the landscape. Real landscapes are more dynamic and can change over longer time scales.

Given the ultimate goal of integrating disease outcomes into land-use planning and management decisions, the model presented here would optimally be run alongside other models in order to estimate tradeoffs between disease risk and ecosystem services that also vary spatially (e.g., see Kennedy et al., 2016). Doing so would allow planned restoration projects to simultaneously optimize over ecosystem services, disease transmission, and other priorities, hopefully helping to avoid a scenario of decreased human health. While data gaps may currently preclude this model’s use directly for decision-making in systems with many hosts and vector species or any understudied systems, its ability to model any number of host and vector species (a strength of the Next Generation Framework generally, which this model draws upon: Schenzle 1984, Anderson and May 1985, Dobson 2004) allows it to be used to predict metrics of disease risk (R_0_ and FOI) in virtually any location where ecosystem services are also of interest or have already been modeled, and extended to land use types beyond urban areas and forest.

It is clear and robust that different infectious diseases will respond differently to land-use changes. It is also highly unlikely that at a landscape scale a given change in tree cover or urban area will lead to an increase or decrease in all relevant infectious diseases. Thus, for any planned restoration project or intervention to combat disease transmission on a changing landscape, it will be paramount to identify which diseases are the most important human health priorities in a given area, as well as which diseases could possibly expand in a changing landscape. Doing so will not only allow public health resources to be targeted proactively as landscapes change, but also allow land-use decision-making to incorporate realistic estimates of the costs and benefits of different scenarios for human health and well-being.

## Acknowledgements

We thank the Mordecai lab for feedback on an early version of the model. EAM was supported by the National Science Foundation (DEB-1518681), the King Center for Global Development, and the Terman Award. EAM and MPK were supported by the National Institute of General Medical Sciences (R35GM133439). MPK was supported by the Natural Capital Project. AJM was supported by the National Science Foundation (DEB-2032276). EAM, LAM, and AJM were supported by the National Science Foundation and the Fogarty International Center (DEB-2011147).

## Competing Interests

The authors declare no competing interests.

## Open Research statement

All data and code used in this study are available in the online supplemental material. Code and data are also hosted at: https://github.com/morgankain/Land-Use_Disease_Model.

## Notes

### Competing Interest Statement

The authors have declared no competing interest.

