## Supplemental Methods and Figures for "Land-use planning for health: Tradeoffs and nonlinearities govern how land-use change impacts vector-borne disease risk"

### Host infection profiles

#### Dengue

Though the role of most non-human hosts involved in sylvatic cycles of dengue transmission remain unknown, dengue is thought to be able to persist in sylvatic cycles; dengue infection and/or antibodies have been detected in a number of mammal species (De Thoisy et al., 2004) including rodents (Thoisy et al., 2009, Cigarroa-Toledo et al., 2016), sloths (Catenacci et al., 2018), bats (Sotomayor-Bonilla et al., 2014), and non-human primates (Vasilakis et al., 2011). Sylvatic cycles are also thought to occasionally begin because of spillover from human-to-human urban transmission (Figueiredo, 2019). However, it remains unclear how competent these species are for transmitting infection to mosquitoes (Cabrera-Romo et al., 2014). For quantitative measures of the transmission capability of host species for dengue, we were only able to find data on human dengue viral load (Duong et al., 2015). Because we failed to find specific pathogen load responses for any other hosts, we simply assumed that non-human hosts are unable to transmit dengue (at least the strains that infect humans) to mosquitoes, and thus we calculated $\mathcal{R}_0$ for dengue starting solely with an infection in a human. We include "other" hosts as sources for mosquito blood meals only. Fitted human dengue viral load data is shown in Figure S1A.

#### Yellow Fever

Non-human primates serve as the primary source of human infection for yellow fever (Kaul et al., 2018, Childs et al., 2019, de Almeida et al., 2019), though yellow fever has also been detected in non-primates (De Thoisy et al., 2004), including rodents (Cigarroa-Toledo et al., 2016) and sloths (Catenacci et al., 2018). However, like for dengue, it remains unclear how competent most of these species are for transmitting infection to mosquitoes (Cabrera-Romo et al., 2014). However, both hamsters (Tesh et al., 2001) and rhesus macaques (Smetana, 1962) have been shown to develop a high enough viral load to infect *Aedes* and *Haemagogus* mosquitoes. Fitted hamsters and macaque profiles are shown in Figure S1B. Here we took the response of rhesus macaques (Smetana, 1962) as the response of non-human primates for the sylvatic transmission component of yellow fever.

For yellow fever we were unable to find quantitative human infection profile data. As, Monath and Vasconcelos (2015) write "Nevertheless there are few data on quantitative viremia levels in humans with yellow fever, and such data would be exceptionally valuable in assessing the potential for vector infection...". However, given the abundance of yellow fever epidemics in human history (e.g. Lee and Moore, 1972, Nasidi et al., 1989, Murphy, 2014), human-to-human transmission is clearly possible and was likely the predominant mode of transmission in historical human epidemics. While vaccination and vector control greatly reduces yellow fever epidemic potential today, the intrinsic ability for an infected human to infect mosquitoes is likely somewhat high. Two additional published comments suggest that human-to-mosquito transmission of yellow fever occurs: 1) Gardner and Ryman (2010) write "During this so-called period of infection, which lasts several days, viremic titers are sufficiently high for transmission to biting mosquitoes" (but provide no citations or data); 2) Vasconcelos and Monath (2016) state that recent epidemics of yellow fever in Africa suggest that *Ae. aegypti* can pick up infection from humans and spread infection to other humans. For these reasons, we assumed that humans mount the same response as that fitted to the species infected in the laboratory studies (Smetana, 1962, Tesh et al., 2001).

#### Malaria

There is an abundance of malaria-causing *Plasmodium* parasite species that tend to be species specific (**?**), though recent work has detected *Plasmodium falciparum* and *Plasmodium vivax* (the two primary agents of human malaria) in multiple species of non-human primates in Colombia (Rondón et al., 2019) and Gabon (Prugnolle et al., 2011). In Vietnam, other work has shown that humans are regularly exposed to primate-specialist *Plasmodium* species, some of which produce disease in humans (Maeno et al., 2015) (though other work finds little human risk of non-human *Plasmodium* specialist species Sundararaman et al. 2013). All in all, it remains unclear how many of these *Plasmodium* species are able to be passed from human to human, and which species are most likely to spill over into humans. For these reasons, we initially cast a wide net in our data collection, taking data (quantitative sampling of pathogen load and/or verbal descriptions of human and non-human pathogen loads) for a variety of *Plasmodium* (Burkot and Graves, 1994, Killeen et al., 2006, Prugnolle et al., 2010, Rayner et al., 2011, Schaer et al., 2013, Bousema et al., 2014, Otto et al.,

2014, Moreira et al., 2015, Vallejo et al., 2016, Martins-Campos et al., 2018). However, because of both large uncertainty in which *Plasmodium* are relevant to humans and our interest in modeling how *Plasmodium* transmitted by *Nyssorhynchus darlingi* varies by land-use, we used data from *Plasmodium vivax* to construct a pathogen load profile for humans and assumed "others" were unable to transmit malaria. A fitted malaria profile for humans is shown in Figure S1C. We appreciate that this malaria pathogen load profile does not represent the long incubation period and serial interval of malaria (Huber et al., 2016). Because we measure one generation of infection for $\mathcal{R}_0$ and $\mathrm{FOI_h}$, what matters is period of time humans are infectious to mosquitoes. While this duration may also be underestimated here, a lack of pathogen load data over time makes it difficult to parameterize a more accurate quantitative infectious profile. Given that we ignore the long incubation period, this profile is not suited for calculating any form of growth rate ("little *r*") or for any multi-generation simulation.

All raw extracted quantitative data are available in the supplemental data file "host_titer.csv".

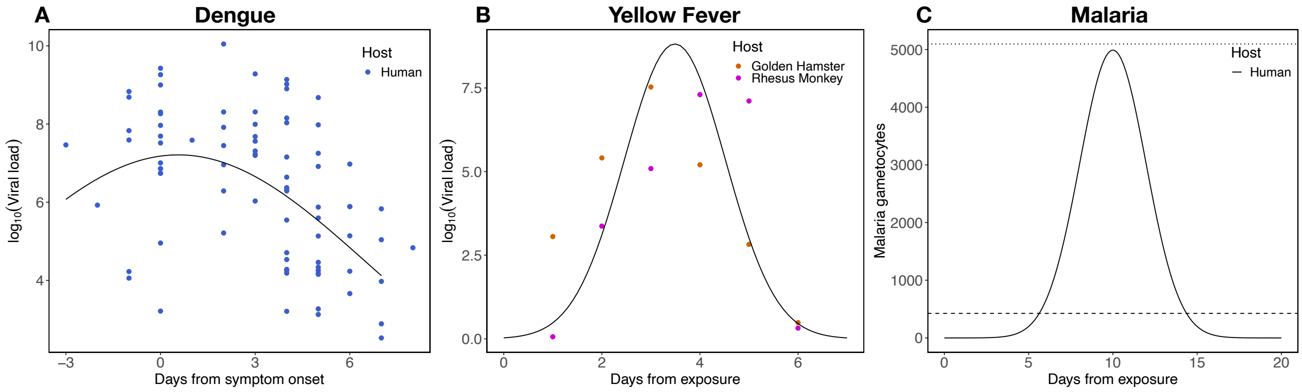


Figure S1: **Host pathogen response profiles.** Linear models with a quadratic term for "day" fit to empirical viral load data from humans infected with dengue (panel A), and golden hamsters and rhesus monkeys infected with yellow fever (panel B). We were unable to find daily malaria gametocyte data for any species, but did find a report of average gametocyte abundance in early and acute phases of malaria infection in humans (Vallejo et al., 2016) (shown as dashed and dotted lines in panel C). Keeping with a quadratic function, we translate these averages into a continuous profile (panel C).

### Mosquito infection and transmission probability

#### *Aedes*

Both *Ae. aegypti* and *Ae. albopictus* are known to be competent for dengue and yellow fever (Mitchell, 1991) (indeed, *Aedes aegypti* was long referred to as the "yellow fever mosquito": Christophers 1960), and both of have been found infected in the field (dengue: De Figueiredo et al. 2010; yellow fever: Fontenille et al. 1997, Possas et al. 2018). Though laboratory infection experiments of both *Ae. aegypti* and *Ae. albopictus* for dengue and yellow fever do exist, papers that report infection probability across dose and transmission probability over time are somewhat rare. Thus, we combined all laboratory experimental data presented from each paper we were able to find reporting *Ae. aegypti* and *Ae. albopictus* infection or transmission probability for yellow fever and dengue (Aitken et al., 1977, Tabachnick et al., 1985, Mitchell et al., 1987, Miller et al., 1989, Johnson et al., 2002, Jupp and Kemp, 2002, Mutebi et al., 2004, Nguyen et al., 2013, Duong et al., 2015, Couto-Lima et al., 2017). *Ae. aegypti* and *Ae. albopictus* are not able to vector malaria.

***Anopheles/Nyssorhynchus***

*Anopheles* spp. are the primary vectors of malaria. Given the variety of *Anopheles* species infected in laboratory experiments and the variety of *Plasmodium* species and strains used in these experiments; we gathered and combined laboratory experimental data from all *Anopheles-Plasmodium* species pairs for which we could find data (Collins et al., 2002, Bharti et al., 2006, Paaijmans et al., 2012, Blanford et al., 2013, Rios-Velásquez et al., 2013, Bousema et al., 2014, Moreno et al., 2014, Churcher et al., 2015, Kiattibutr et al., 2017). However, in an effort to tailor the model to specifically the transmission of malaria in South America by *Ny. darlingi*, we restricted our estimates of infection to that of *Ny. darlingi* (we found no data for *Ny. darlingi* specific transmission probabilities). *Anopheles/Nyssorhynchus* spp. are not able to transmit dengue or yellow fever.

***Haemagogus***

Finally, the primary vectors for the sylvatic cycle (non-human primates-to-mosquitoes-to-primates) of yellow fever are the *Haemagogus* and *Sabethes* genera of mosquitoes. Here we combined these two genera into a single sylvatic specialist mosquito *type*; we refer to this aggregate simply as *Haemagogus* because we mostly used *Haemagogus* parameters (though we did include a small number of *Sabethes* parameters). We obtained data on *Haemagogus* mosquitoes previously collected for Childs et al. (2019) (data available at: `https://github.com/marissachilds/YellowFeverSpillover/tree/master/data`).

**Models**

We modeled mosquito infection as virus detected in the abdomen and mosquito transmission as virus detected in either salivary glands or heads in order to utilize as much as data as possible (many old studies grind up entire mosquito heads while newer studies extract virus from salivary glands only). Given the ubiquity of defining infection as virus in the abdomen on (or near) day 14, we fit with day as a fixed effect but estimated infection probability for use in our transmission model using day 14. Both the infection and transmission model assume that the probability of infection/transmission reaches one after some dose or time period, which is not the case for many arbovirus infections. While it would be prudent to model a variable maximum infection and transmission probability by species, we lacked the data to do so for most mosquito-pathogen pairs. Thus, we used simple logistic regression which models probability using an asymptote of one. For the purposes of mosquito transmission, for computational reasons (a massive increase in array size), we did not track the dose (pathogen load) every mosquito receives when biting the host depending on the day each mosquito bit the host. Instead we assumed mosquitoes pick up the average dose from each host they bite (for transmission probability).

Fitted mosquito infection probability—the proportion of mosquitoes exposed to a given pathogen load (in an infection experiment the dose given to the mosquito, and in our model the pathogen load within a host that the mosquito bites: see Figure S1) that end up infected (have detectable pathogen in their bodies), which is used in the host-to-mosquito arm of the transmission cycle—is shown for dengue and yellow fever in Figure S2A and for malaria in Panel B. Unfortunately, the literature search conducted by Childs et al. (2019) and our own additional literature search failed to find papers that recorded mosquito infection. Most of these papers are from the 1930's through 1970's and used actual feeding experiments of mosquitoes on live hosts instead of extracting virus from mosquitoes, which is a methodology that is unable to quantify infection probability at the level of the individual mosquito. Given this lack of data, we simply assumed that mosquito infection probability for *Haemagogus* is identical to that of *Aedes albopictus*.

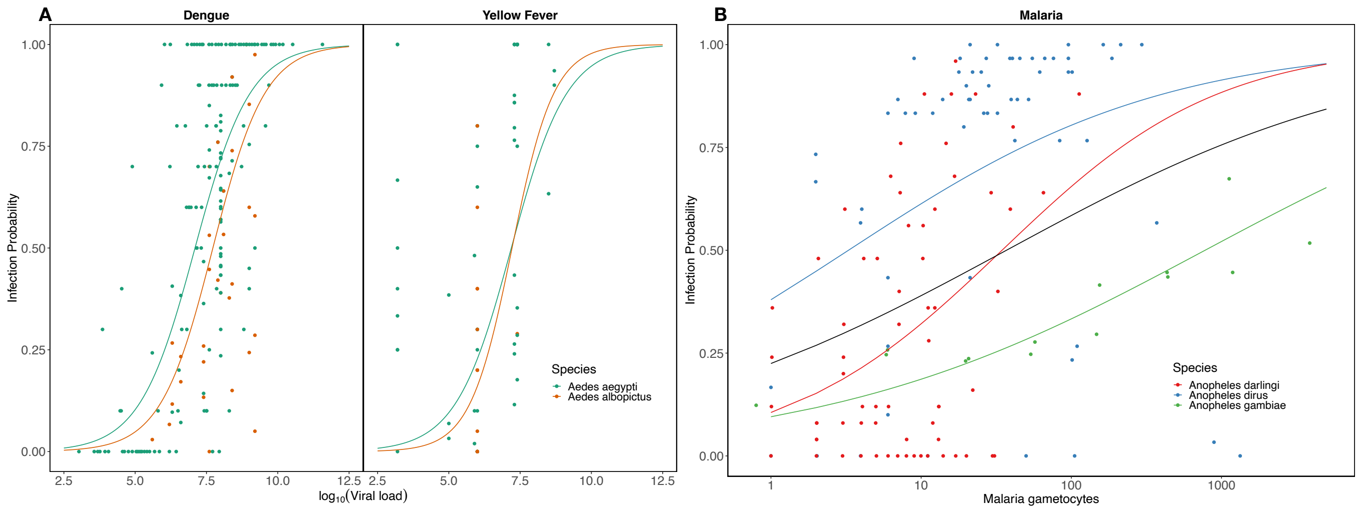


Figure S2: **Mosquito infection probability.** Logistic regression fits to experimental data on mosquito infection probability for dengue and yellow fever (panel A), and malaria (panel B).

Fitted mosquito transmission probability—the proportion of mosquitoes exposed to a given pathogen load that are able to transmit the pathogen (have detectable pathogen in their salivary glands), which is used in the mosquito-to-host arm of the transmission cycle—is shown in Figure S3A-C. Estimated mosquito transmission probability irrespective of mosquito survival (an aggregate of the curves in Panels A-C) is shown in Figure S3D, and mosquito survival and transmission probability weighted by survival are shown in Figure S3E-F, respectively. Given both the spread of the data and the similarity in transmission probability of dengue and yellow fever by the two *Aedes* species, we modeled these probabilities together (Figure S3A). Because the experimental data that we were able to find for malaria transmission were obtained across a broad temperature range, we included temperature in the model for *Anopheles* transmission of malaria (we used *An. stephensi* for this model because of available data and thus assumed the response of *Ny. darlingi* was identical to that of *stephensi*. To predict malaria transmission on the landscape we used the median of the temperatures (23ºC).

Given poor data from very old papers (1930s-1970s) on *Haemagogus* transmission of yellow fever, we relied on a subset of the transmission data collected for Childs et al. (2019). All experiments did not use virus in salivary glands but instead let pools (usually) of mosquitoes (rarely individual mosquitoes) feed on an infected host and then feed, after some number of days for incubation, on a susceptible host. The host was then monitored to determine if it became infected. The issue with these data are that there is no way to tell how many of the mosquitoes got infected, just that at least one mosquito was infected and then was able to transmit. These data do help give a general picture of extrinsic incubation period, but do not provide the granularity that we needed here (proportion of mosquitoes transmitting over time). However, a small subset of these experiments did track individual mosquitoes; we restricted our model to using these data. One additional issue of note with these subset data is the use of variable doses, strains and other laboratory techniques (e.g., temperature and housing conditions). Further, many of these covaraties are simply not reported. Thus, while we were able to fit a model to literature data, the data is noisy and the fit is poor (Figure S3C).

All raw extracted quantitative data for dengue, yellow fever, and malaria are available in the two supplemental data files “mosquito_to_host_transmission.csv” and “mosquito_to_host_transmission.csv”

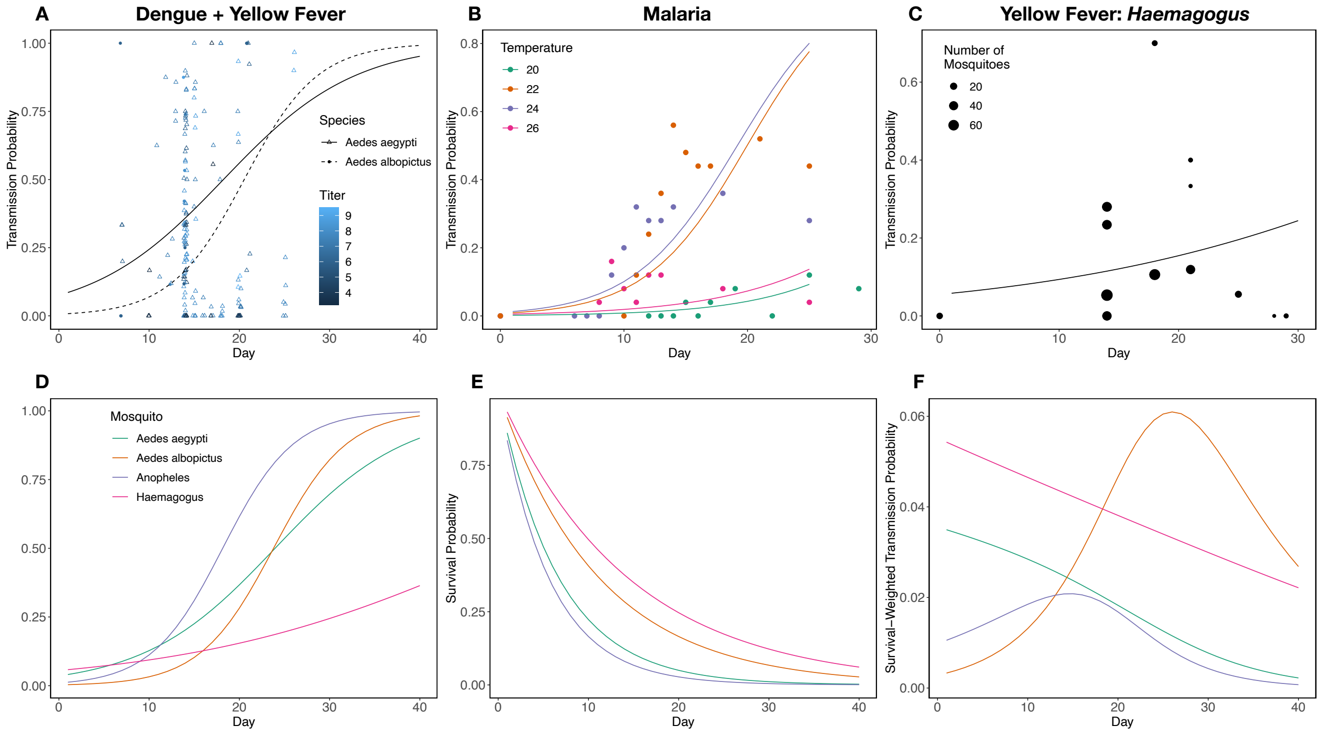


Figure S3: **Mosquito transmission probability.** Mosquito traits that affect transmission: transmission probability and survival probability. Points and triangles in panel A show raw data for the probability *Aedes aegypti* and *Aedes albopictus* have viral load in their salivary glands or head across days and infecting dose (which is the way "transmission" is experimentally measured). Points and triangles are a mix of dengue and yellow fever infections; solid and dotted lines are predicted transmission probability at a dose of 7 $\log_{10}$ viral load (for visual purposes we just chose one infectious dose). Colored points in panel B show raw data for the probability *Anopheles stephensi* have detectable malaria parasites in their salivary glands across days for temperatures ranging from 20-26°C; colored lines show model predictions. The points in panel C show raw data on the proportion of *Haemagogus* mosquitoes transmitting yellow fever; the line shows the predicted mean response. Single curves in panel C show the estimates used for the landscape transmission model; transmission estimated at 7 $\log_{10}$ viral load for *Aedes* and at 23°C for *Anopheles*. Survival probabilities, calculated using exponential decay, of these four mosquito species are shown in panel D. Transmission probability weighted by survival probability is shown in panel E.

### Mosquito biting behavior

For all mosquito species we gathered data on the origin of their blood meals from field-based blood meal analyses of wild caught mosquitoes. In total we obtained blood meal information from: Charlwood et al. (1995), Tandon and Ray (2000), Kiszewski et al. (2004), Alencar et al. (2005), Ponlawat and Harrington (2005), Richards et al. (2006), Zimmerman et al. (2006), Alencar et al. (2008), Delatte et al. (2010), Kamgang et al. (2012), Moreno et al. (2017). All raw extracted quantitative data for all mosquito species are available in the supplemental data file "feeding_preference.csv".

### Mosquito biting rate

Mosquito biting rate is notoriously variable, possibly varying as much as an order of magnitude among locations (Sallum et al., 2019), and across temperature (Mordecai et al., 2013). Here we assume a moderate biting rate for all three mosquito species using the average of the Breire-modeled biting rate between 20 and 40 for *Ae. aegypti* and *Anopheles pseudopunctipennis* from (Mordecai et al., 2013) as a basis, which returns 0.37 and 0.55 bites per day for these two species respectively. We assume a biting rate of 0.37 for *Ae. albopictus* as well, but increase *Ae. aegypti* by 1.5-fold because of their ability to bite multiple times per *Haemagogus* mosquitoes we used the inverse of their estimated gonotrophic cycle length from (Dégallier et al., 1998) because we were unable to find any direct estimates of daily biting rate.

### Host abundance

For the transmission of both dengue and malaria "other" hosts includes all non-human species including primates. Only for yellow fever do we separate "others" into primates and "others". For the yellow fever model, in the absence of better data, we simply assumed the abundance of "others" and primates were equal. While this is somewhat of an arbitrary assumption, the favored feeding location of *Haemagogus* mosquitoes in the mid-canopy (Dégallier et al., 1998) will increase the ratio of primates to all other species. We stress that the relative density of "others" and primates will affect $\mathcal{R}_0$ values when starting with either a human infection or a primate infection. Thus, our $rzero$ values for yellow fever should be interpreted with caution.

### Mosquito abundance

We combine data on the locations of wild caught mosquitoes with long standing knowledge of the preferred habitats of *Aedes aegypti*, *Aedes albopictus*, *Ny. darlingi*, and *Haemagogus* spp. mosquitoes (Braks et al.,

2003, Scott and Morrison, 2010, Sarfraz et al., 2012, 2014, de Moura Rodrigues et al., 2015, Mucci et al., 2015, de Camargo-Neves et al., 2005, Lin et al., 2016, Tátila-Ferreira et al., 2017, Pereira dos Santos et al., 2018, Delatorre et al., 2019, Koyoc-Cardeña et al., 2019, Hendy et al., 2020, Silva et al., 2020) to establish absolute estimates of mosquito abundance on our simulated landscapes. Specifically, we use a *highly flexible*, multi-parameter exponential function to sculpt a quantitative relationship between mosquito density and landscape characteristics: urbanization, human density, tree cover, and, "other"species density. We note that: 1) while this function is set up *like* a regression model that has individual coefficients associated with continuous covariates, we do not actually statistically *fit* this function to any data, instead simply "parameterize" this function to translate qualitative knowledge into a quantitative relationship; 2) the advantage of such a flexible function is that it can model nearly any desired relationship; however, this flexibility also increases the danger that the model's predictions are overly dependent on the relationships we assumed here, which are choices and not model output fitted to quantitative data; 3) this is a key place where our model could be improved. For example, a mosquito species distribution model with forest cover and human population density fit to mosquito capture data could be used to obtain regression coefficients, which could then be used to predict mosquito densities on simulated landscapes (given that the regression model's predictors span their range in the simulation).

The function we used is as follows:

$$\begin{aligned} \rho &= (1 + e^{-(a + b\cdot\omega + c\cdot\omega^2 + d\cdot\delta)})^{-1} \\ \theta &= \frac{\rho}{max(\rho)} \lambda \psi^{\psi_{exp}} \end{aligned} \tag{1}$$

where $\theta$ gives the absolute density of a given mosquito species in a landscape cell; the precursor $\rho$ gives the relationship between landscape features and relative mosquito density before being scaled to absolute density. The intermediate product $\rho$ is calculated using four coefficients multiplied by three unique continuous covariates: $a$ is a constant intercept term; $b$ and $c$ scale a linear and quadratic relationship between mosquito abundance and a primary landscape feature $\omega$ (for *Ae. aegypti* urbanization; *Ae. albopictus*, *Ny. darlingi*, and *Haemagogus*: tree cover); $d$ scales the relationship between mosquito abundance and a secondary landscape feature $\delta$ (*Ny. darlingi*: variance in tree cover in an area around the focal pixel defined by their flight radius; unused for *Aedes* spp. and *Haemagogus* spp.). To calculate absolute density ($\theta$), we scale $\rho$ to [zero, one] and multiply it by $\lambda\psi^{\psi_{exp}}$, where $\psi$ is absolute host abundance (as defined as a weighted mean of human and "other" host species, weighted by mosquito biting preference), and $\lambda$ and $\psi_{exp}$ are linear and exponential scaling factors, respectively, that provide the relationship between host abundance the mosquito population sizes such a host abundance can support. We assume a default value of 1 for $\psi_{exp}$ for *Ae. aegypti* to model a linear dependence, but explore a range of parameters to examine sub- and supra-linear dependencies (Romeo-Aznar et al., 2018).

In Figure S4, Figure S5, Figure S6, Figure S7, we show the output of this function for *Ae. aegypit*, *Ae. albopictus*, *Anopheles*, and *Haemagogus* abundance, respectively, on one of our simulated landscapes.

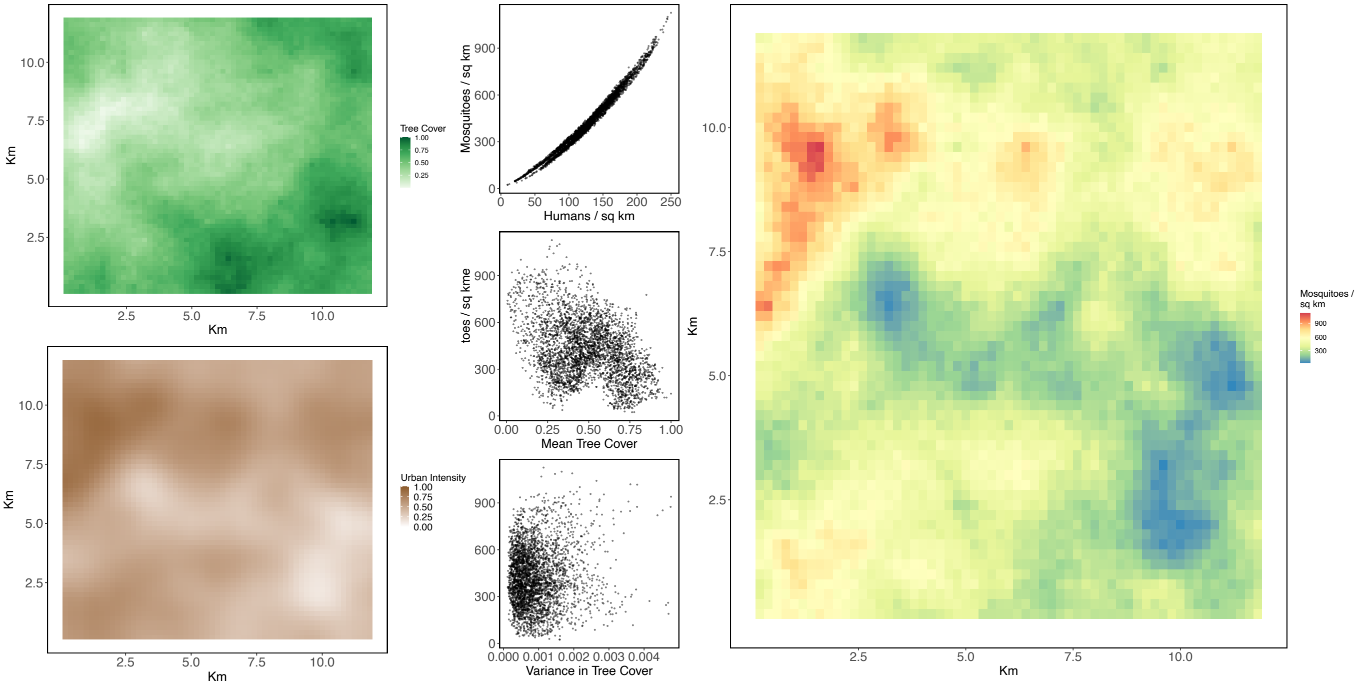


Figure S4: ***Aedes aegypti* population density on a sample simulated landscape with spatial autocorrelation of 0.48**. The left column shows simulated tree cover and human population density for this sample landscape. The center column shows estimated mosquito density as a function of human population density, mean tree cover, and variance in tree cover in $\pm 3$ pixels in each cardinal direction. The matrix on the right shows estimated mosquito population density on the landscape. *Aedes aegypti* population density is driven almost entirely by human population density. Parameters used to model *Aedes aegypti* abundance were as follows: a = 0.5, b = 0.75, c = 0, d = 0, $\psi_{exp} = 1$, $\psi$ = simulated human and "other" abundance weighted by *Aedes aegypti* feeding preference, $\delta$ = not used for *Aedes aegypti*, $\lambda$ = 5.

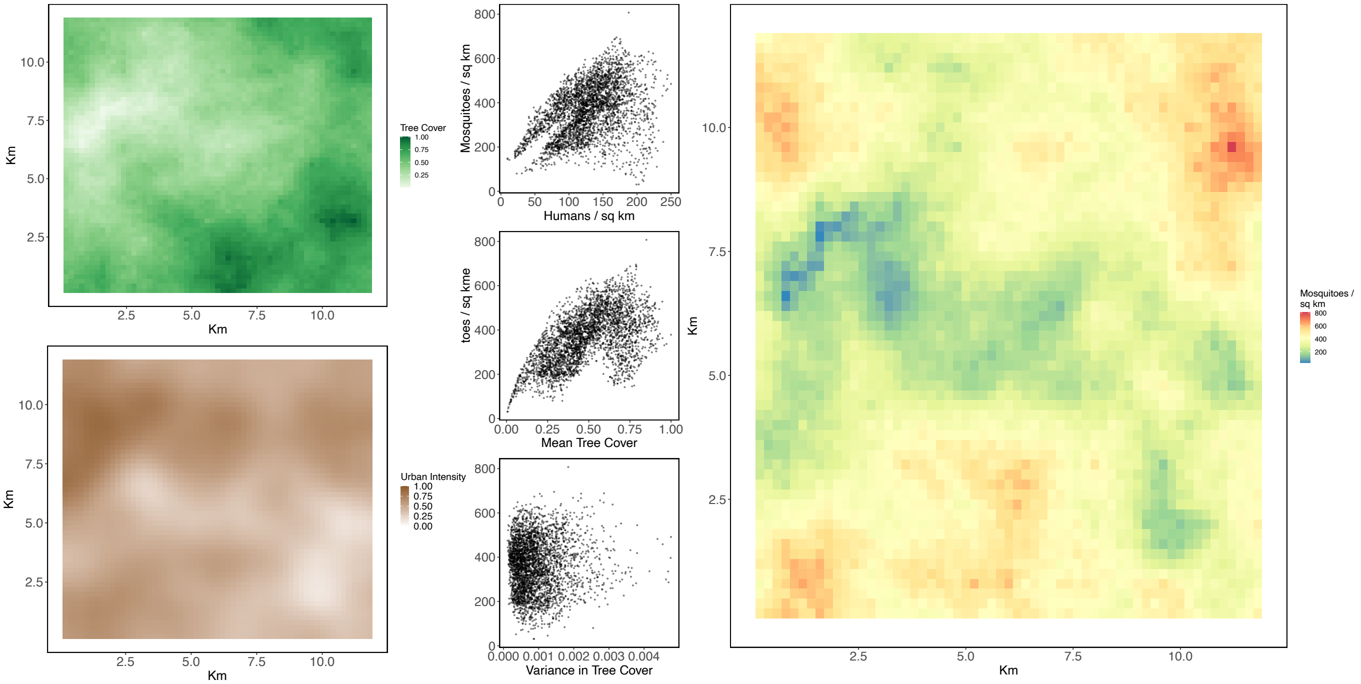


Figure S5: ***Aedes albopictus*** **population density on a sample simulated landscape with spatial autocorrelation of 0.48**. The left column shows simulated tree cover and human population density for this sample landscape. The center column shows estimated mosquito density as a function of human population density, mean tree cover, and variance in tree cover in ±3 pixels in each cardinal direction. The matrix on the right shows estimated mosquito population density on the landscape. *Aedes albopictus* population density is driven by a combination of human population density and mean tree cover. Parameters used to model *Aedes albopictus* abundance were as follows: a = 0.5, b = 0.75, c = 0, d = 0, $\psi_{exp} = 1$, $\psi =$ simulated human and "other" abundance weighted by *Aedes albopictus* feeding preference, $\delta$ = not used for *Aedes albopictus*, $\lambda = 5$.

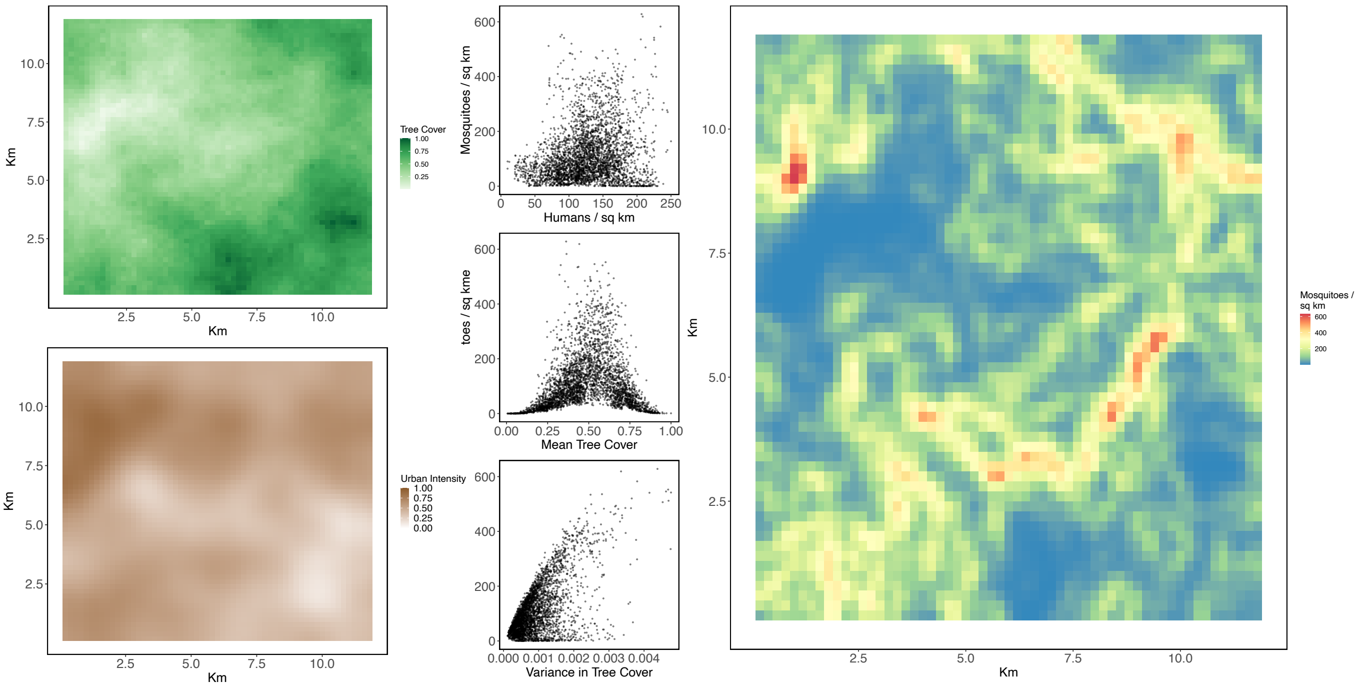


Figure S6: ***Ny. darlingi* population density on a sample simulated landscape with spatial auto-correlation of 0.48**. The left column shows simulated tree cover and human population density for this sample landscape. The center column shows estimated mosquito density as a function of human population density, mean tree cover, and variance in tree cover in $\pm 3$ pixels in each cardinal direction. The matrix on the right shows estimated mosquito population density on the landscape. *Ny. darlingi* population density is driven primarily by the combination of mean tree cover and variance in tree cover. Parameters used to model *Ny. darlingi* abundance were as follows: a = 1, b = 0, c = -1, d = 1, $\psi_{exp} = 1$, $\psi =$ simulated human and "other" abundance weighted by *Ny. darlingi* feeding preference, $\delta$ = log(variance in tree cover $\cdot$ 50), $\lambda$ = 5.

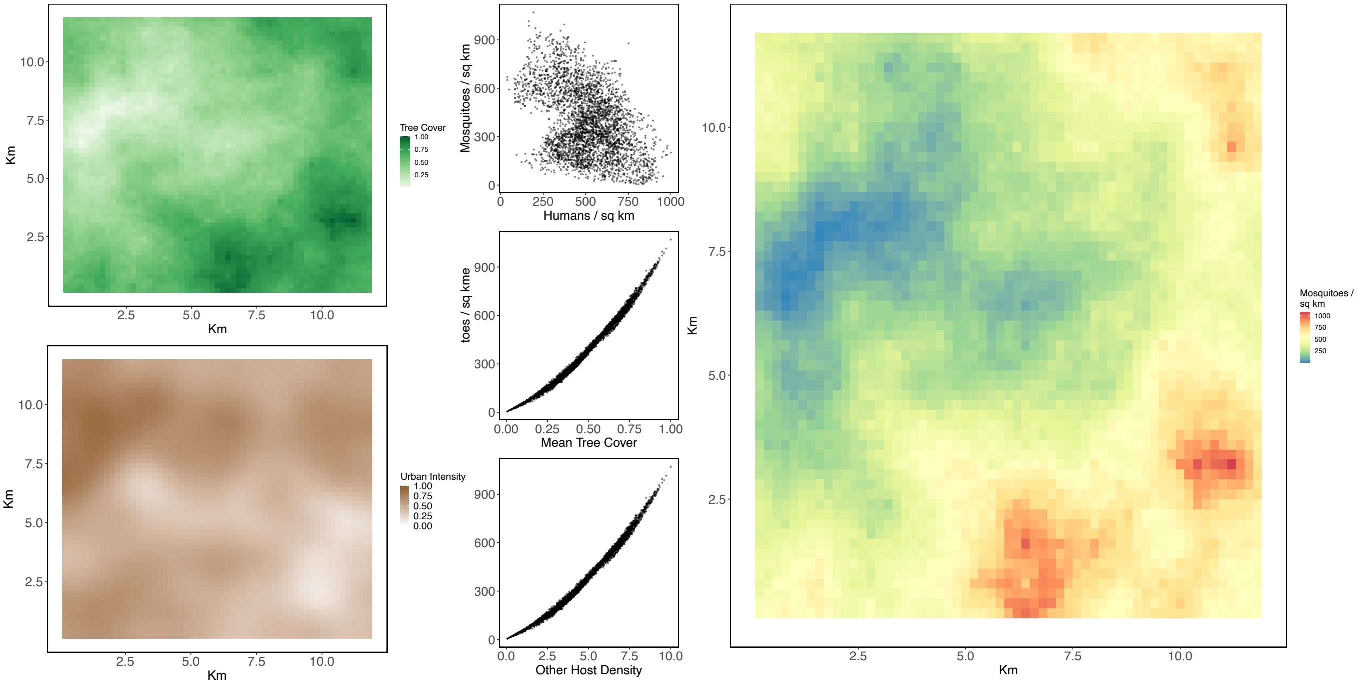


Figure S7: ***Haemagogus* spp. population density on a sample simulated landscape with spatial autocorrelation of 0.48**. The left column shows simulated tree cover and human population density for this sample landscape. The center column shows estimated mosquito density as a function of human population density, mean tree cover, and variance in tree cover in ±3 pixels in each cardinal direction. The matrix on the right shows estimated mosquito population density on the landscape. *Haemagogus* are generally thought to be forest specialists that breed in tree holes (Tátila-Ferreira et al., 2017, Silva et al., 2020), though they are also found in patchy forest and human-disturbed habitats (Medeiros-Sousa et al., 2015, Mucci et al., 2015, Delatorre et al., 2019). Here we model them as a forest specialist, but in the model do not weight their flight by tree cover, thus allowing them to readily disperse into urban areas. Specifically, we model their abundance using most of the same parameters as for *Aedes albopictus*: a = 0.5, b = 0.75, c = 0, d = 0, $\psi_{exp} = 1$, $\delta$ = not used for *Haemagogus*, $\lambda$ = 5, though $\psi$ = simulated human and "other" abundance weighted by *Haemagogus* feeding preference.

### Simulated landscapes

Example simulated landscapes for the four values of spatial auto-correlation used to show results in the main text are shown in Figure S8.

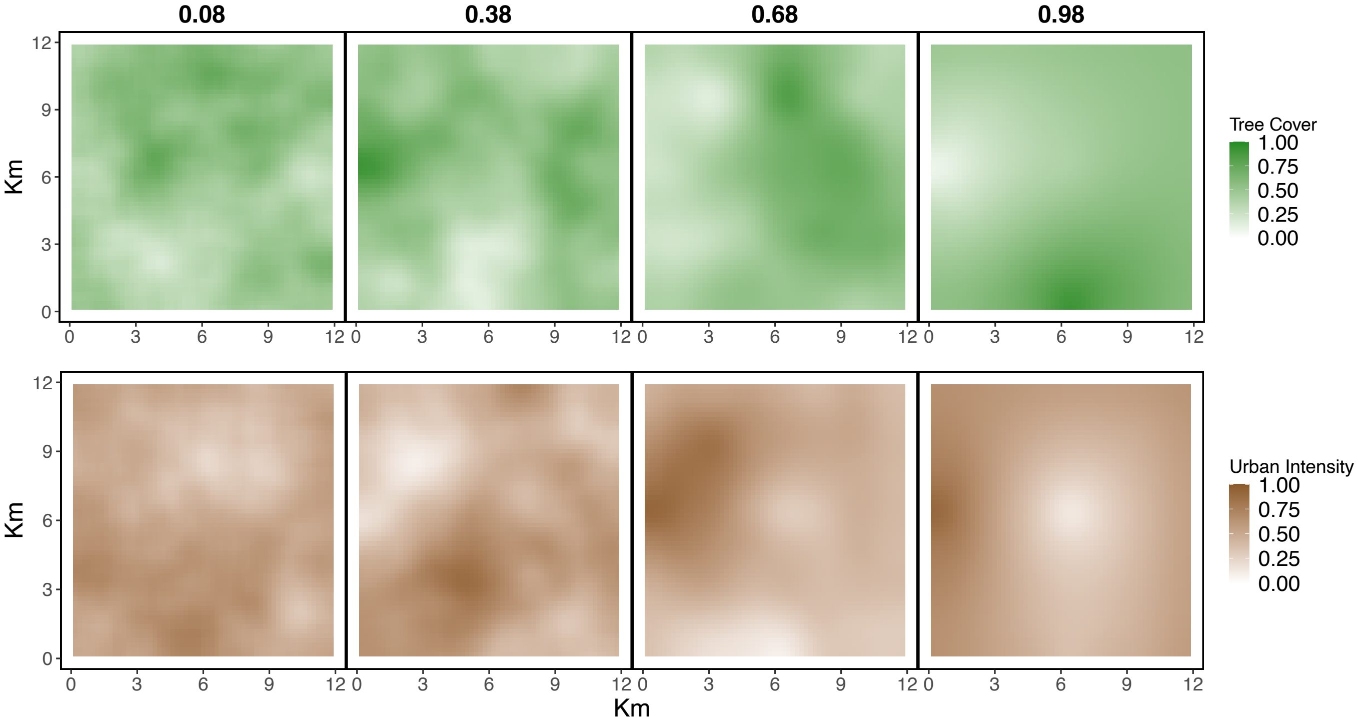


Figure S8: **Simulated tree cover and urbanization for landscapes across four values of spatial auto-correlation.** As spatial auto-correlation increases, tree cover and urbanization are constrained to be more contiguous. Given the negative correlation between these landscape features, the greater the spatial auto-correlation the closer to the "land-sparing" extreme the landscapes become.

### Empirical landscape

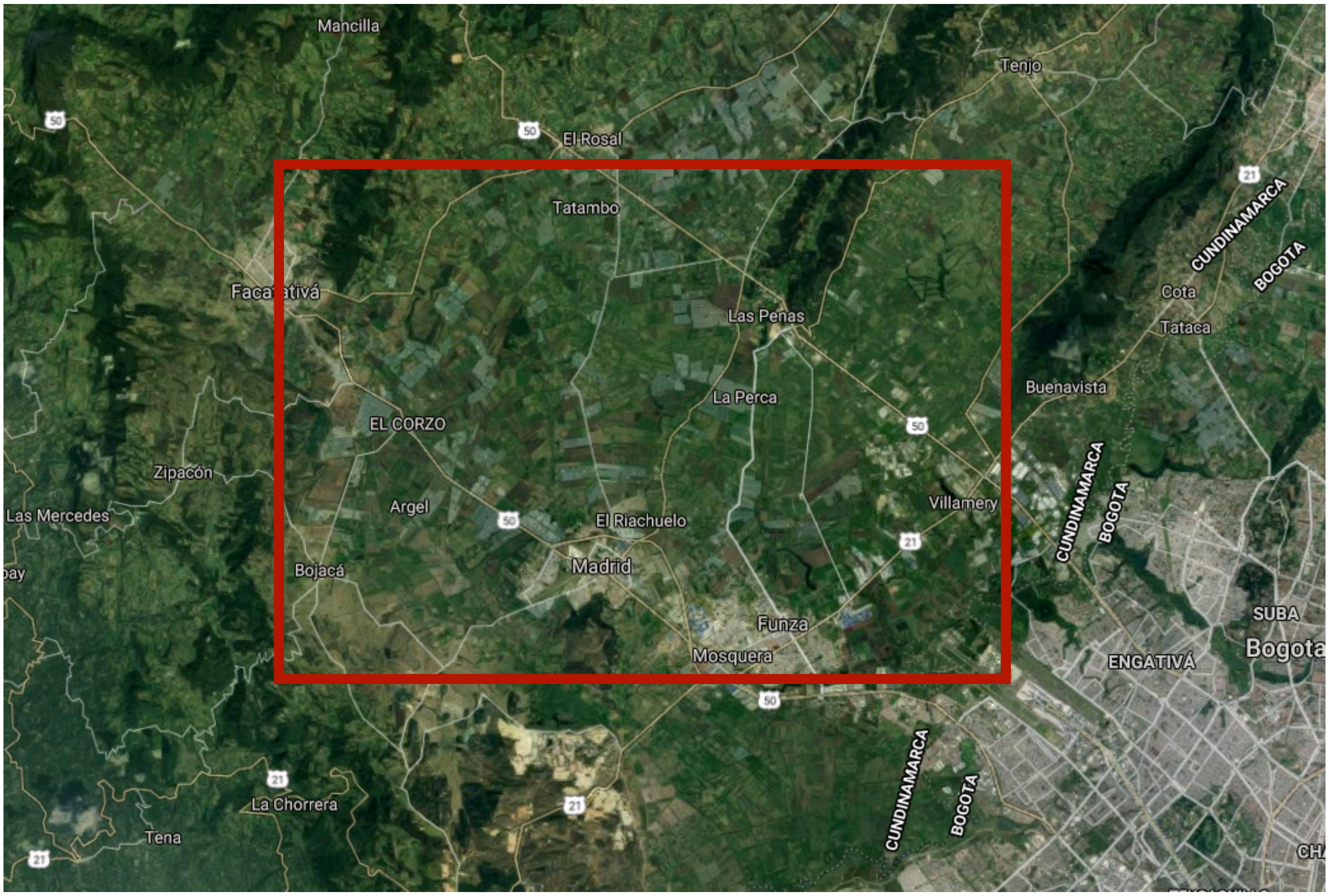


Figure S9: **Northwestern outskirts of Bogota, Columbia and the surrounding peri-urban area, farmland, and forest cover used as the landscape for our empirical example.** The full image was obtained from Google Maps satellite view; the internal red rectangle is the area covered in our analysis.

### Supplement: Additional Results

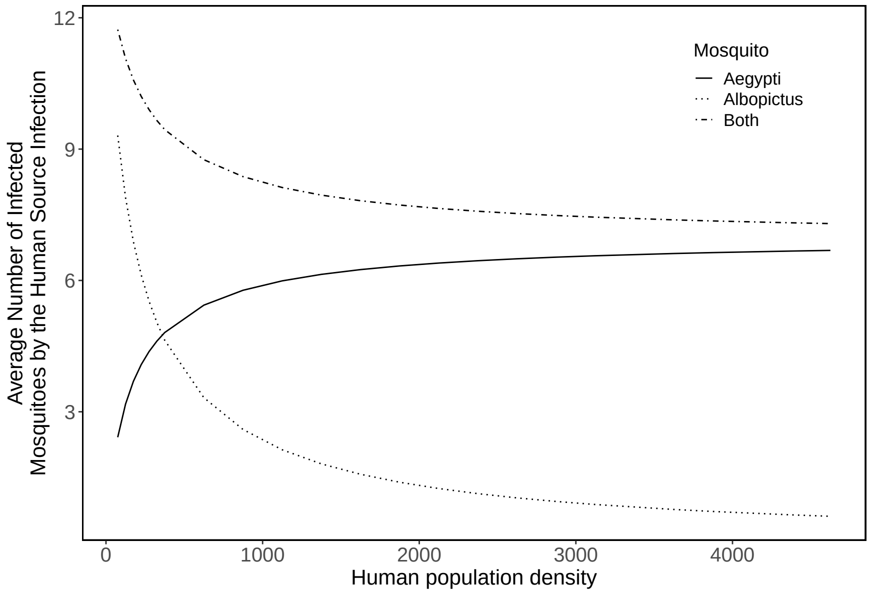


Figure S10: **The number of *Aedes aegypti* and *Aedes albopictus* infections resulting from a source human infection, averaged across a source infection in every cell on a landscape**. As human population density increases, the number of *Aedes albopictus* that get infected from the single source infection decreases. This occurs because *Ae. albopictus* abundance is assumed to be independent of human population density, which means that the single source human dengue infection becomes "lost" in a sea of susceptible humans to the population of blood feeding *Ae. albopictus*. Because the number of infected *Ae. albopictus* decreases with increasing human population density, fewer second-generation human infections are generated by *Ae. albopictus*; see Figure 3).

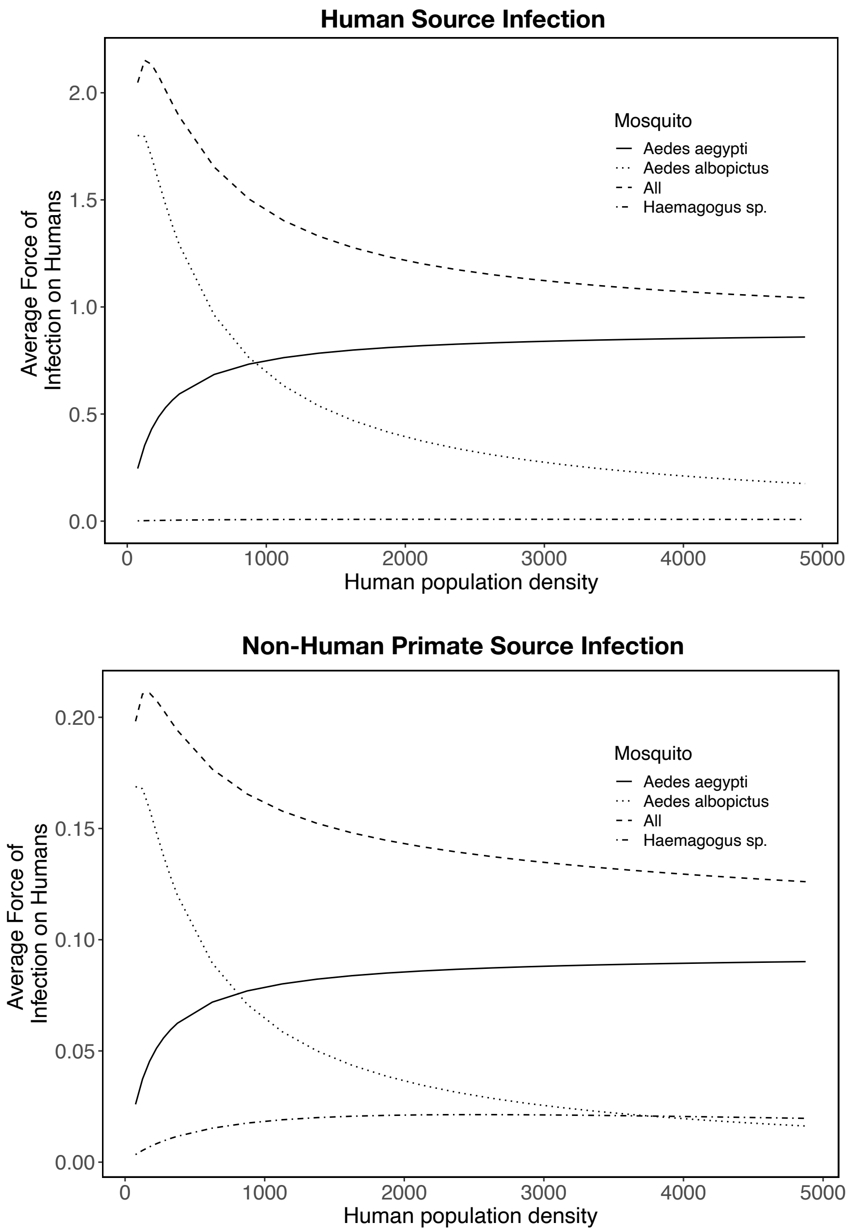


Figure S11: **Average yellow fever $FOI_h$ on simulated landscapes that vary in absolute human population density, partitioned into the relative contributions made by *Aedes aegypti*, *Aedes albopictus*, and *Haemagogus* spp.**. Top panel shows $FOI_h$ from a human source infection and the bottom panel shows $FOI_h$ from a non-human primate source infection. Total $FOI_h$ is the sum of the contributions made by *Aedes aegypti*, *Aedes albopictus*, and *Haemagogus* spp. mosquitoes. The overall $FOI_h$ from a human infection is much larger than from a primate infection because of the high competence of *Aedes aegypti* and *Aedes albopictus* for transmission and the high biting affinity of these mosquitoes for humans. These results were calculated on a landscape with a spatial auto-correlation of 0.78.

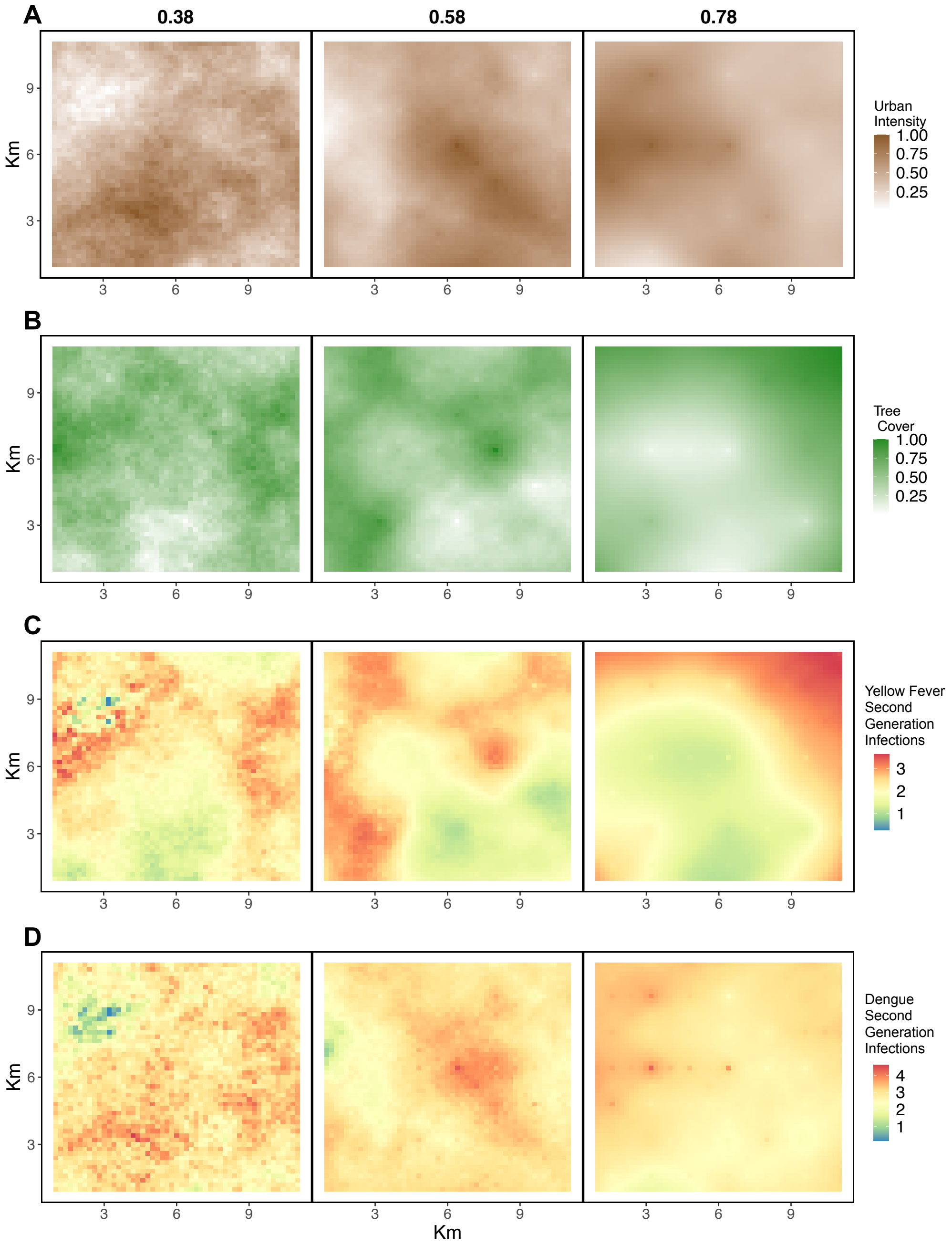


Figure S12: **Relationship among dengue and yellow fever $FOI_h$ on three simulated landscapes with medium-low (0.38) medium (0.58) or medium-high (0.98) spatial autocorrelation for urban area and tree cover.** Panels A and B show urban intensity and tree cover respectively, while panels C and D show yellow fever $FOI_h$ and dengue $FOI_h$. All results pictured here are for landscapes with average human population density of 250 people per sq.km.

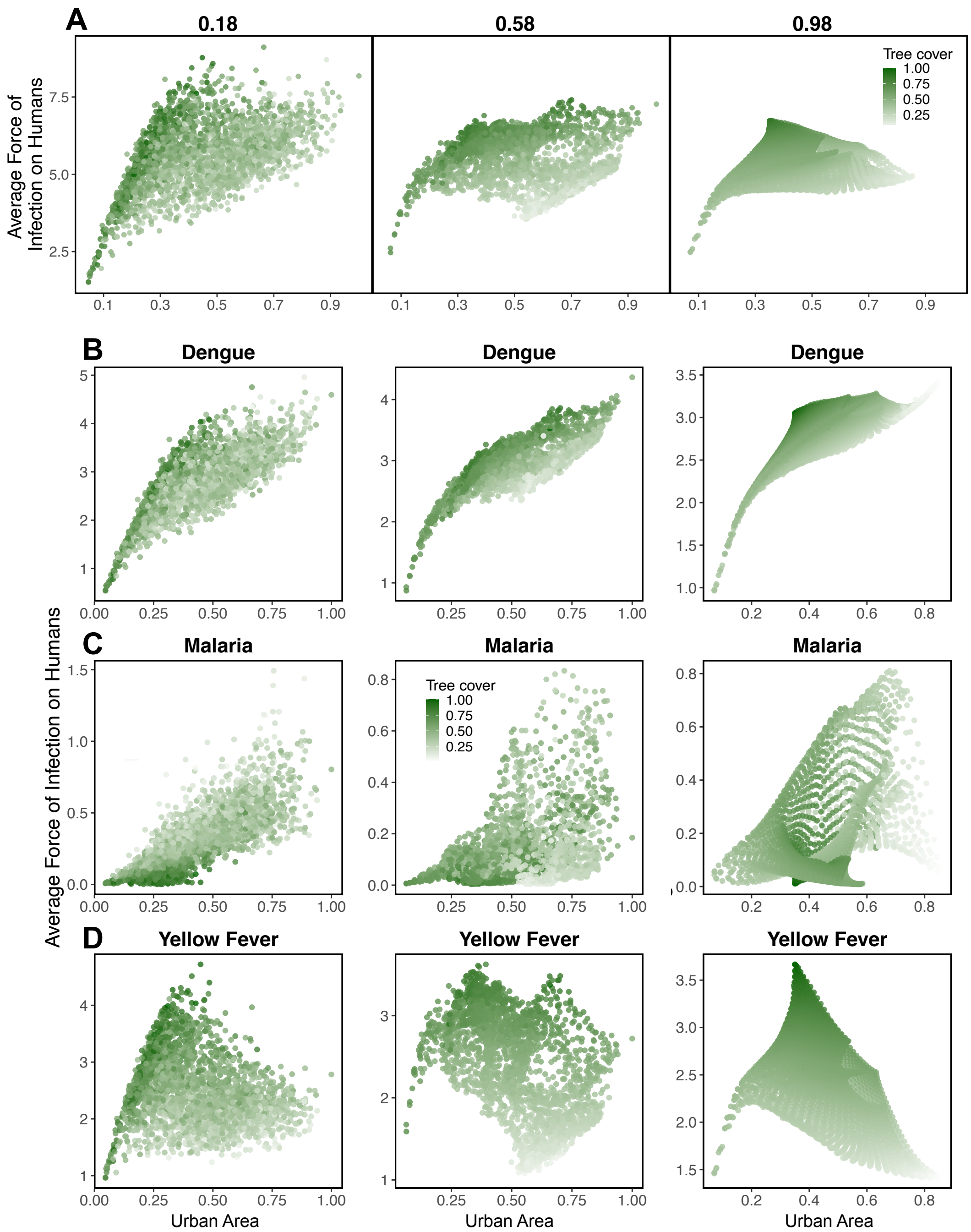


Figure S13: **Relationship among the $FOI_h$ of each disease on three simulated landscapes with low (0.18) medium (0.58) or high (0.98) spatial autocorrelation for urban area and tree cover.** Panel A shows the total $FOI_h$ summed across all three diseases. Plots in rows B-D show the relationship between the $FOI_h$ of each disease and urban area and tree cover; each point shows an estimate from a single landscape cell. All results pictured here are for landscapes with average human population density of 250 people per sq.km.

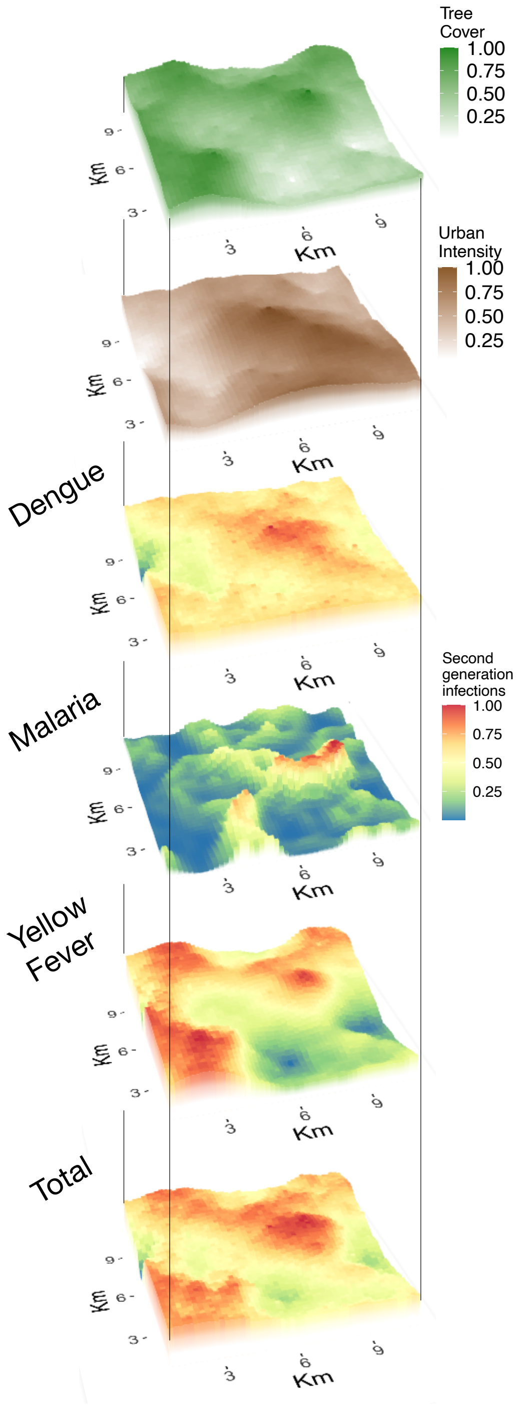


Figure S14: **Simulated landscape features and estimated $FOI_h$ of each disease, aligned and stacked to aid visualization of the overlap of features with high-risk and low-risk regions of the landscape.** This simulated landscape is the same as that with medium spatial auto-correlation among landscape features (0.58) shown in Figure 5, which has an average human population density of 250 people per sq.km.

Instead of the somewhat contrived assumption of infections arising on each landscape cell, disease risk hotspots can alternatively be modeled assuming that only a *single* infection of each disease were to appear somewhere on the landscape. In this case, the probability of emergence of each disease on each cell must be written as a function of landscape features, for which many sensible alternative relationships exist. Here we calculated $\mathrm{FOI_h}$ in this way assuming that dengue emergence probability was directly proportional to human population abundance, while yellow fever and malaria emergence probabilities were directly proportional to primate abundance and *Ny. darlingi* abundance, respectively. With these emergence probability weights, the spatial patterns of disease risk on the Bogotá landscapes (Figure S15) are similar to the $\mathrm{FOI_h}$ pictured in Figure 8. However, the spatial regions of high risk are smaller for all diseases on all landscapes when disease risk is calculated using a single infection; dengue risk is more constrained to areas of high human population density, malaria risk is maximized in the regions with the most variation in forest cover, and yellow fever risk is constrained to regions with high tree cover (Figure S15).

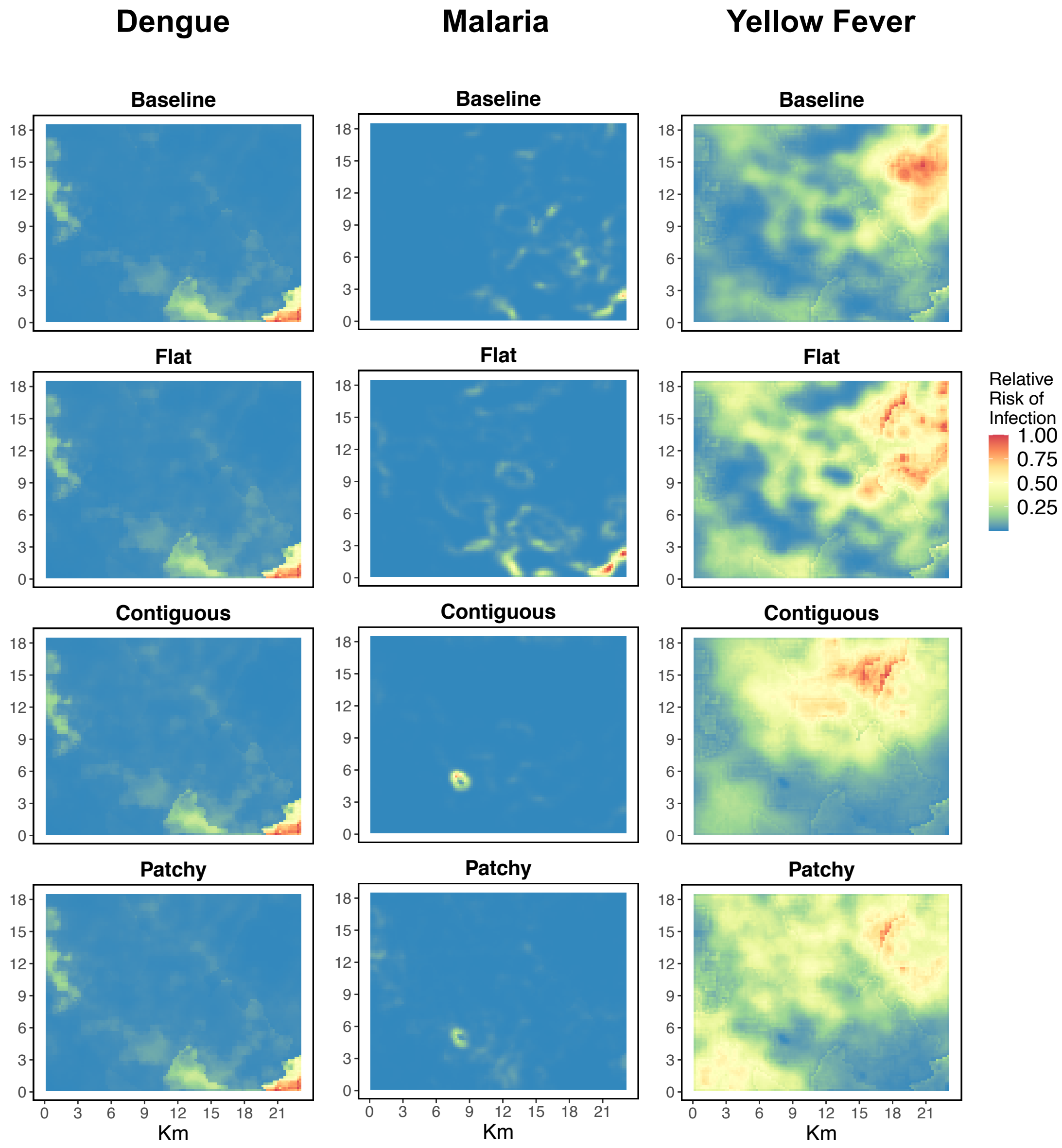


Figure S15: **Estimated relative $FOI_h$ of each disease on a 23km x 18km landscape to the northwest of Bogotá, Columbia for three potential scenarios of reforestation, assuming the a single infection arises somewhere on the landscape**. Maps of the estimated relative $FOI_h$ of dengue, malaria, and yellow fever are shown for "Baseline" and for each reforestation scenario: "Flat" increased tree cover evenly across the landscape (which serves as a null-model), "Congtiguous" simulated the planting of a single large patch of forest (e.g., a regional conservation effort), and "Patchy" simulated the planting of many small patches (e.g., subsidies to individual farms to replant trees). For all scenarios the average tree cover on the landscape is simulated to be brought up from 0.14 (as measured by LAI; see supplemental methods) at baseline to 0.50.
